# Substrate-interacting pore loops of two ATPase subunits determine the degradation efficiency of the 26S proteasome

**DOI:** 10.1101/2023.12.14.571752

**Authors:** Erika M. López-Alfonzo, Ayush Saurabh, Sahar Zarafshan, Steve Pressé, Andreas Martin

## Abstract

The 26S proteasome is the major eukaryotic protease responsible for protein quality control, proteostasis, and the modulation of numerous vital processes through the degradation of regulatory proteins. Commitment to degradation occurs when conserved pore loops in the proteasomal heterohexameric ATPase motor engage the flexible initiation region of a polyubiquitinated protein substrate for subsequent mechanical unfolding and translocation into a proteolytic chamber. Here, we used *in vitro* biochemical and single-molecule FRET-based assays with mutant reconstituted 26S proteasomes from yeast to characterize how the pore-1 loops of individual ATPase subunits in the AAA+ motor contribute to the different steps of substrate degradation and affect the proteasome conformational dynamics. We found that the pore-1 loop of the Rpt6 ATPase subunit plays particularly important roles in substrate capture, engagement, and unfolding, while the pore-1 loop of the Rpt4 ATPase is critical for providing sufficient grip for substrate unraveling and maintaining a processing-competent state of the proteasome. Interestingly, these pore-1-loop contributions correlate with their positions in the spiral-staircase arrangements of ATPase subunits in the substrate-free and substrate-degrading proteasome, providing new insights into the mechanisms of substrate processing by the 26S proteasome and related hexameric ATPase motors.

## Introduction

Protein degradation is essential for cellular homeostasis, cell division, differentiation, quality control, and the regulation of numerous other vital processes. The Ubiquitin Proteasome System (UPS) acts as the major pathway for energy-dependent protein degradation in eukaryotic cells, with the 26S proteasome as the final component recognizing and degrading targeted protein substrates ^1^. This 2.5 MDa macromolecular machine is composed of the cylindrical 20S core particle (CP) and the 19S regulatory particle (RP) that caps the 20S CP on one or both ends (Fig. 1A). The 20S CP contains proteolytic active sites in an internal chamber for polypeptide cleavage, whereas the 19S RP is responsible for the recognition of ubiquitinated substrates, their deubiquitination, mechanical unfolding, and translocation into the proteolytic chamber. The 19S RP consists of the lid and base subcomplexes. The lid subcomplex with its nine subunits binds to one side of the base and core, and contains the essential Zn^2+^-dependent deubiquitinase Rpn11 ^2–4^. The base is made up from ten subunits, including three main ubiquitin receptors (Rpn1, Rpn10, and Rpn13) ^5–8^, the scaffolding subunit Rpn2, and six distinct ATPases (Rpt1-Rpt6) of the AAA+ (<u>A</u>TPases <u>A</u>ssociated with diverse cellular <u>A</u>ctivities) family that form the heterohexameric motor of the proteasome. These ATPase subunits, arranged in the order Rpt1, Rpt2, Rpt6, Rpt3, Rpt4, and Rpt5 (as seen from the top of the proteasome)^9^, share a common domain architecture that includes a N-terminal coiled-coil, a small domain with an oligonucleotide/oligosaccharide-binding (OB) fold that in the hexamer forms a N-terminal domain ring (N-ring), and a C-terminal ATPase domain consisting of a large and a small AAA+ subdomain that assemble into the ATPase motor ring ^10^. This ATPase ring docks on top of the 20S CP and induces opening of the axial gate for substrate access to the proteolytic chamber ^11^. Each ATPase domain contains conserved motifs for ATP binding (Walker A) and ATP hydrolysis (Walker B), as well as a pair of pore loops, called the pore-1 and pore-2 loop, that project into the central channel of the hexameric ring ^12^. The pore-1 loops sterically contact the substrate polypeptide through conserved tyrosine and lysine residues for ATP-hydrolysis-driven substrate unfolding and translocation ^11,13–16^.

**Fig. 1.**
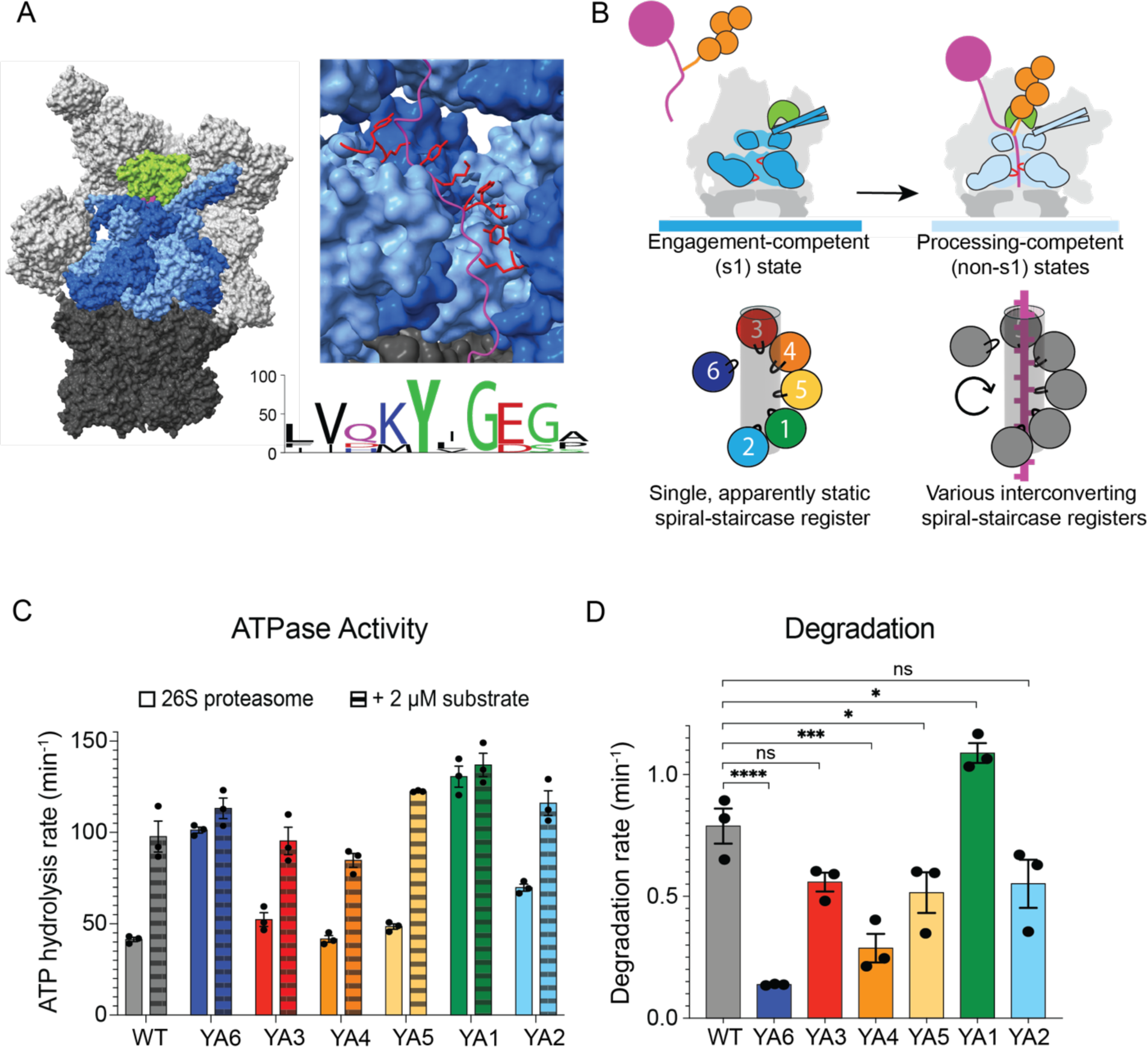
Pore-1 loop mutations have differential effects on proteasome activities. **A)** Left: Structure of the 26S proteasome during substrate processing (PDB ID: 6EF3), with the 20S core particle in dark grey, the lid subcomplex and the none-ATPase subunits of the base subcomplex in light grey, the Rpn11 deubiquitinase subunit of the lid in lime green, the six Rpt ATPase subunits of the base alternating in two shades of blue, and the substrate polypeptide inside the central channel in magenta. Top right: Side view of the base subcomplex with two ATPase subunits, Rpt3 and Rpt4, removed for a better view of the central channel with an engaged substrate chain (magenta). Pore-1 loops (shown in red cartoon representation) with a conserved Lys-Tyr motif (or Met-Tyr for Rpt5) project from each subunit into the central channel and surround the substrate in a spiral staircase conformation. Bottom right: Sequence logo for the pore-1 loops of the yeast Rpt subunits shows the strong conservation of the Lys-Tyr motif. **B)** The pore-1 loops form a specific spiral-staircase arrangement in the engagement-competent s1 state (dark blue motor), with Rpt3 at the top, Rpt2 at the bottom, and Rpt6 at the intermediate “seam” position. Upon substrate engagement, the regulatory particle shifts to processing-competent non-s1 states (light blue motor) with various spiral-staircase registers that may represent intermediates of a hand-over-hand mechanisms in which subunits move from the bottom via the seam position to the top of the staircase. **C)** Rates for ATPase activity of reconstituted proteasomes in the absence (solid bars) and presence of 2 μM ubiquitinated titin I27^V15P^ substrate (dashed bars). Technical replicates (N = 3) are plotted with error bars representing the SEM. **D)** Rates for the degradation of FAM-labeled ubiquitinated titin I27^V15P^ substrate by reconstituted proteasomes under single turnover conditions, measured by changes in fluorescence polarization. (N = 3 technical replicates, error bars represent the SEM). Statistical significance was calculated using a one-way ANOVA test: ****p<0.0001; ***p<0.001;**p<0.01; *p<0.05; ns, p>0.05.

To be degraded by the 26S proteasome, a substrate must be modified with several ubiquitins, usually in the form of a polyubiquitin chain, and contain an unstructured initiation region for engagement by the ATPase motor ^17–20^. Substrate-attached ubiquitin chains are recognized by one or more ubiquitin receptors of the proteasome, followed by insertion of the unstructured initiation region into the central channel of the ATPase hexamer. Cryo electron-microscopy (cryo-EM) studies showed that in the absence of substrate the proteasome primarily resides in an engagement-competent conformation, or s1 state, in which the entrance to the central channel is accessible ^21–29^. Upon substrate insertion and engagement by the motor, the proteasome undergoes a conformational change from s1 to non-s1, processing-competent states. During this transition, the lid shifts and rotates relative to the base, and the Rpn11 deubiquitinase becomes coaxially aligned above the central channel, such that it partially obstructs the entrance, and the engaged substrate needs to be translocated through a small gap, which facilitates efficient ubiquitin removal. ATP-dependent translocation of the substrate by the AAA+ motor then drives mechanical unfolding, co-translocational deubiquitination by Rpn11 ^30^, and transfer of the unstructured polypeptide into the CP’s internal degradation chamber for proteolytic cleavage.

Cryo-EM structures of the proteasome in the absence and presence of substrate revealed that the six Rpts and their pore-1 loops form different ordered spiral-staircase arrangements (Fig. 1B) ^21–29,31–37^. The engagement-competent s1 state is characterized by a single staircase register, with Rpt3 at the top, Rpt2 at the bottom, and Rpt6 at a “seam” position, located between the top and bottom subunits. Structures of the substrate-engaged proteasome in processing-competent ^31,37^, non-s1 states showed various spiral-staircase registers with different subunits at the top, bottom, and seam positions, different nucleotide occupancies. In these non-s1 states, three to five subunits toward the top of the staircase were found engaged through their pore-1 loops with the substrate polypeptide in the central channel. These observations led to a sequential hand-over-hand translocation model ^31^, in which the penultimate subunit in the staircase hydrolyzes ATP, causing the neighboring ADP-bound bottom subunit to disengage from the substrate, move as the “seam” subunit to the top of the staircase, exchange ADP for ATP, and re-engage with the substrate. The bottom or “seam” subunits in the staircase arrangements of s1- and non-s1-states are therefore expected to be the next Rpt to bind and pull on the substrate, and for the s1 state this step may be critical for stable substrate engagement and inducing the conformational switch to non-s1 processing conformations. However, these models are solely based on structural snapshots of substrate-free and substrate-engaged proteasomes, and detailed biochemical and biophysical analyses are necessary to test these models and elucidate the principles of ATP-hydrolysis-driven substrate translocation.

Through ensemble and single-molecule Fluorescence Resonance Energy Transfer (smFRET) assays ^17,38^, we previously showed that switching from the engagement-competent s1 to the processing-competent non-s1 states upon substrate insertion into the ATPase motor is a key step for the commitment to degradation and that mutations shifting the conformational equilibrium toward non-s1 states inhibit substrate engagement and processing. For instance, mutations of the VTENKIF motif in the Rpn5 lid subunit that break s1-state-specific contacts between the lid and base subcomplexes or mutation of the Walker-B motif in Rpt6 that traps this subunit in the permanent ATP-bound state both shift the conformational equilibrium toward non-s1 states, and thereby inhibit substrate insertion and degradation ^34,39^.

Previous biochemical studies also attempted to determine the importance of individual pore-1 loops for substrate processing. When pore-1 loop tyrosines were individually replaced with glycine in *S. cerevisiae*, the Rpt4 mutant showed the highest accumulation of ubiquitinated conjugates in whole-cell lysates ^40^. In contrast, when substituting the pore-1 loop tyrosine with an alanine (referred to as YA mutation) in yeast cells, mutant Rpt1 (YA1) and Rpt6 (YA6) exhibited the most significant growth defects at different temperatures ^13^. Our own *in vitro* biochemical studies of YA mutants found differential defects in substrate degradation, ATP hydrolysis, and CP gate opening, with YA1 being the most unaffected mutant ^11^.

These variable defects observed for pore-1 loop mutants *in vivo* and *in vitro* do not allow deriving a conclusive mechanistic model for ATP-dependent substrate processing and the functional contributions of individual Rpts, and a more detailed characterization of mutant effects on the various degradation steps and on the proteasome conformational dynamics is needed.

Here, we used smFRET-based assays to conduct a comprehensive mechanistic dissection of the pore-1 loop functional asymmetries in the proteasomal AAA+ ATPase motor. Our experiments revealed that the ATPase subunits located at the bottom or “seam” positions in the Rpt spiral-staircase arrangements of substrate-free and substrate-degrading proteasomes are particularly important for substrate capture, robust unfolding without release, processive translocation, and controlling the proteasome conformational transitions. These findings point to a mechanism for substrate engagement and mechanical unfolding in which the intrinsic asymmetry of the proteasomal motor and particular spiral-staircase arrangements of Rpts play critical roles at different stages of ATP-dependent degradation.

## Results

### Pore-1 loop mutations cause differential degradation defects

To understand how the individual pore-1 loops of Rpt1 – Rpt6 contribute to substrate processing, we recombinantly expressed *S. cerevisiae* base subcomplexes with single Tyr to Ala (YA) pore-1-loop mutations in *E. coli* (Supp. Fig.1-1A) and *in vitro* reconstituted them after purification with recombinantly expressed *S.c.* lid and endogenous 20S CP purified from *S.c.* to form 26S proteasomes. Using recombinant systems for the lid and base subcomplexes not only gives us the advantage of introducing mutations that would otherwise be detrimental to proteasome activity in yeast, but also enables the incorporation of unnatural amino acids for site-specific labeling with fluorescent dyes to perform smFRET-based measurements of substrate processing and proteasome conformational changes ^11,17,38^. As a substrate, we employed a previously characterized non-fluorescent model protein that can be labeled with fluorescent dyes for anisotropy and FRET-based measurements, and is thus well suited for degradation experiments in bulk and at the single-molecule level ^17,38^. This substrate consists of a titin I27 domain with or without a destabilizing V15P mutation (titin I27^V15P^ or titin I27), followed by a 24 amino acid linker that contains a single lysine for enzymatic ubiquitin-chain attachment, and a 35 amino acid unstructured initiation region with a cysteine for labeling with a fluorescent dye (Supp. Fig.1-1A).

To test the functional defects of pore-1-loop mutant proteasomes, we first measured their ATP-hydrolysis and substrate-degradation activities. It was previously shown that addition of substrate to wild-type proteasomes causes a 2-fold stimulation of basal ATPase activity ^11^, likely linked to the conformational switch from the engagement-competent s1 to processing-competent non-s1 states upon substrate engagement. Using a NADH-coupled assay to measure ATP hydrolysis, we observed the expected 2-fold stimulation for reconstituted wild-type proteasomes, with the basal ATPase rate of 41 ± 1 min^-1^ increasing to a rate of 98 ± 8 min^-1^ in the presence of 2 μM ubiquitinated titin I27^V15P^ (Fig. 1C, Supp. Fig.1-2A). The YA3, YA4, YA5, and YA2-mutant proteasomes showed overall similar basal ATPase rates and stimulations by substrate of 1.7 – 2.5 fold. In contrast, YA6 and YA1 proteasomes had strongly elevated basal ATPase rates of 101 ± 2 and 131 ± 6 min^-1^, respectively (Fig. 1C), that did not significantly increase upon substrate addition. An earlier study reported strongly increased ATPase activity for the YA1 mutant purified from yeast ^13^, yet this phenotype was not previously described for YA6.

To test degradation defects of the YA-mutant proteasomes, we measured their kinetics for the single turnover of the titin I27^V15P^ substrate that was labeled with a N-terminal FAM peptide for degradation readout by a decrease in fluorescence polarization (Fig. 1D). If all six pore-1 loops in the substrate-engaged, non-s1 state proteasome contributed equally to unfolding and translocation, as proposed by regular hand-over-hand translocation and sequential ATP-hydrolysis models, one would expect similar defects for all YA mutants. Although we observed that the YA3, YA5, and YA2 mutations had comparable effects (Fig. 1D, Supp. Fig.1-2B,C), YA6 and YA4 proteasomes exhibited much more decreased degradation activities of 20% and 35%, respectively. Interestingly, the YA1 mutant, which similar to YA6 showed an increased basal ATPase rate with no further stimulation upon substrate addition, degraded the model substrate at 135% of the wild-type rate. YA1-mutant proteasomes can thus use their stimulated ATPase rate for faster substrate degradation, whereas a considerable fraction of hydrolysis events for the YA6 mutant are futile. Our bulk degradation data are consistent with previous *in vivo* studies indicating a particular importance of Rpt4, Rpt6, and Rpt1 for substrate processing ^13,40^. However, these results do not provide mechanistic insights on how individual pore-1 loops contribute to the different substrate-processing steps and how the proteasome may utilize its structural asymmetries for efficient degradation. We therefore further investigated the functional defects of pore-1-loop mutant proteasomes, with a particular focus on Rpt4 and Rpt6.

### Pore-1 loops of Rpt4 and Rpt6 influence the proteasome conformational dynamics

Given the strong effects of the YA4 and YA6 mutations on ATP-hydrolysis and substrate-degradation rates, we turned to our previously developed smFRET-based assay that allows monitoring the conformational states of the 26S proteasome and the investigation of YA-mutant effects ^38^. In this assay, the lid subunit Rpn9 is labeled with the FRET-donor fluorophore LD555 at an azido-phenylalanine (AzF) replacing Phe2, and the ATPase subunit Rpt5 is labeled with the FRET-acceptor fluorophore LD655 at an AzF substituted for Gln49 (Fig. 2A). The proteasome conformational switch from the s1 to non-s1 states leads to a > 30 Å decrease in the distance between the fluorophore-attachment points and can therefore be observed by changes in apparent FRET efficiency (app. FRET eff.) ^17^. Reconstituted proteasomes were immobilized on a microscope coverslip through biotin-neutravidin interactions as previously described ^38^, and changes in the donor and acceptor fluorescence intensities were monitored by Total Internal Reflection Fluorescence (TIRF) microscopy (Supp. Fig. 2A). Compared to our previously published studies, we used an upgraded system with an increased signal-to-noise ratio and a faster sampling rate of 20 Hz, and we therefore first re-examined the conformational dynamics of the wild-type proteasome.

**Fig. 2.**
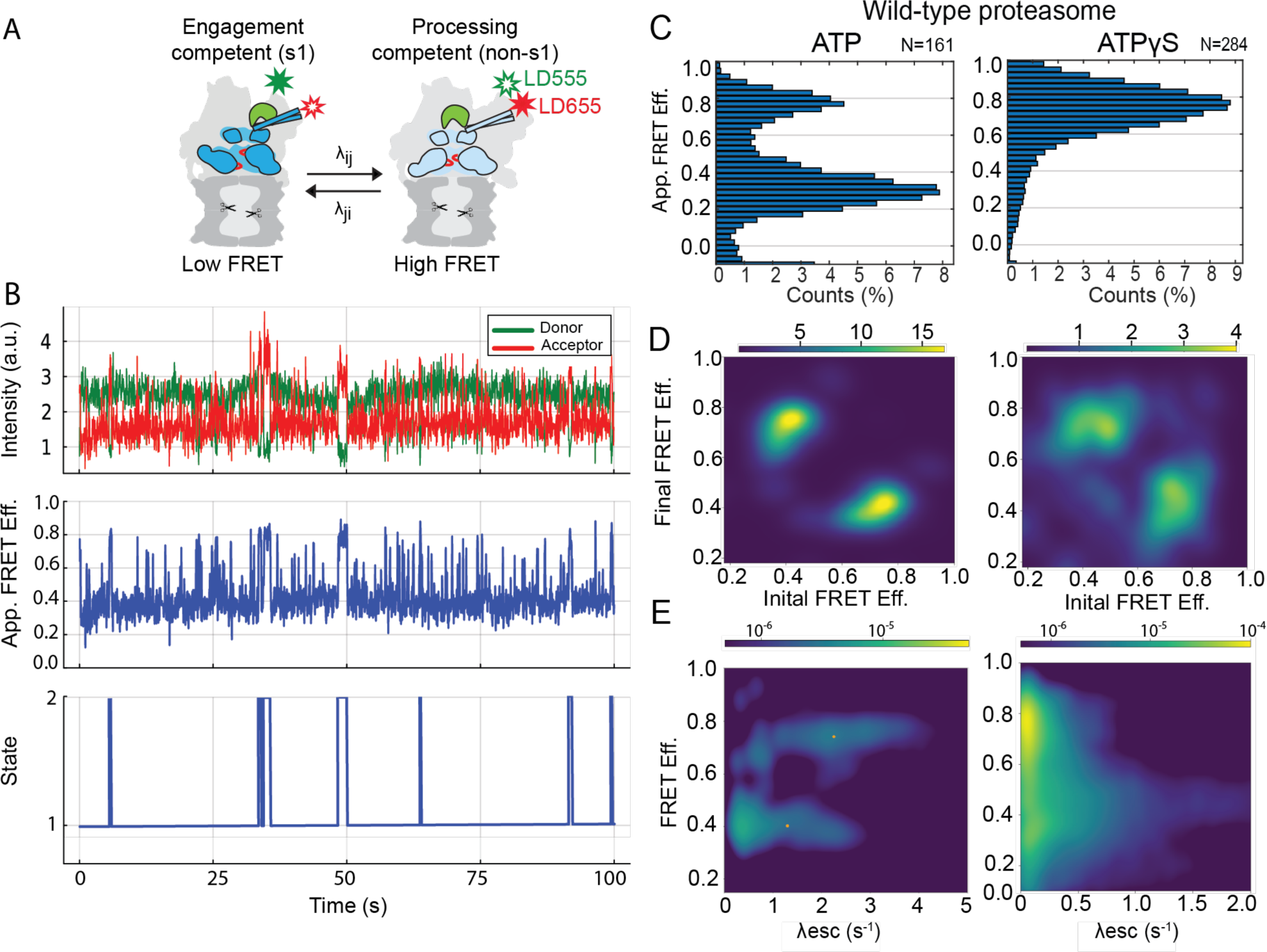
Conformational dynamics of the 26S proteasome in the absence of substrate. **A)** FRET-based assay to monitor the proteasome conformational transitions between the engagement-competent s1 and processing-competent non-s1 states. Proteasomes were labeled with a FRET-donor dye (LD555, green star) on the lid subunit Rpn9 and a FRET-acceptor (LD655, red) on the base ATPase subunit Rpt5. **B)** Representative traces for the single-molecule fluorescence measurement of the wild-type 26S proteasomes in ATP by TIRF microscopy. The top panel shows the fluorescence intensities for the FRET donor (green) and acceptor (red), the middle panel shows the apparent FRET efficiency, and the bottom panel illustrates the estimated most-probable state trajectory. **C)** Histograms of apparent FRET efficiencies for wild-type 26S proteasome in ATP (left, N = 161) or ATPγS (right, N = 284). **D)** Transition density plots for wild-type 26S proteasome in ATP (left) or ATPγS (right). A scale color map (side bar) indicates the density transitions within the sampled states. **E)** Bivariate probability distributions showing the escape rates (λesc) for each FRET efficiency of wild-type 26S proteasomes in ATP (left) or ATPγS (right). To distinguish probability regions, a logarithmic scale color map was used (top bar), indicating the probability of proteasomes to assume a specific FRET efficiency based on all the states sampled.

To calculate the conformational transition rates for proteasomes in the absence and presence of substrate in a more unbiased manner, we utilized Hidden Markov Model - Markov Chain Monte Carlo (HMM-MCMC) sampling techniques, which implement a Bayesian non-parametric framework (Supp. Fig. 2B) that does not pre-determine variables like the number of proteasome states, kinetic rates, and apparent FRET efficiencies. With each smFRET trace acquired (Fig. 2B, top and middle panels), the HMM-MCMC algorithm explores hundreds of possible state trajectories for estimating the most probable one (Fig. 2B, bottom panel). After determining the most probable trajectory, all explored trajectories are pooled to generate transition-density plots and bivariate probability distributions. The transition density plots provide information about the direction of FRET-state transitions and hence the conformational switching of proteasome molecules (Fig. 2D). The bivariate probability distributions allow a visualization of the calculated rates for the escape from each FRET efficiency (λ^esc^ in s^-1^, Fig. 2E) and of the probabilities that proteasomes assume a specific FRET efficiency based on the states sampled (Fig. 2E, color coded bar). To focus on the proteasome conformational transitions, fluorescence traces showing static complexes or events of photobleaching were excluded. In addition, the overall FRET-state distributions of acquired traces were analyzed in apparent FRET efficiency histograms (Fig. 2C).

For wild-type proteasomes in ATP, we observed most particles in the low-FRET, engagement-competent s1 state, with a smaller population sampling high-FRET processing-competent non-s1 states (Fig. 2C, 2D). Based on the bivariate probability distributions, wild-type proteasomes had a mean rate of 1.3 s^-1^ for the escape from the low-FRET-efficiency state (corresponding the s1 - to - non-s1 transition). In contrast, the mean rate for escape out of the high-FRET-efficiency state (corresponding the non-s1 - to - s1 transition) was 2.25 s^-1^. This is ∼ 2-fold lower than the value we previously reported, likely due to the new HMM-MCMC analysis that attempts to incorporate photon shot noise, camera noise, spectral crosstalk, and background emission in a physically accurate manner using camera calibration frames in order to avoid overfitting. Our measurements also revealed the expected behavior for proteasomes in the presence of the non-hydrolysable ATP analog ATPγS that stabilizes the non-s1, high-FRET states (Fig. 2C, D, E).

After establishing our HMM-MCMC data-analysis method for the wild-type proteasome, we collected and analyzed traces for all YA mutants in the presence of ATP and the absence of substrate (Fig. 3B,C, Supp. Fig. 3-1, 3-2, 3-3). YA1, YA2, YA3, YA4, and YA5-mutant proteasomes showed FRET-efficiency distributions similar to wild type (Fig. 3F, Supp. Fig. 3-2), and the degradation-deficient YA4 mutant had also conformational escape rates similar to wild-type (Fig. 3G and 3I, Supp. Table 1), indicating that these pore loops play no major role for the intrinsic rate of the proteasome conformational switching. In contrast, the YA6-mutant proteasome exhibited increased sampling of the processing-competent non-s1 states, as observed in the apparent FRET efficiency histogram (Fig. 3E). This was confirmed by an increased mean s1–to– non-s1 escape rate of 1.57 s^-1^, with a distribution elongated to rates beyond 4 s^-1^ (Fig. 3H). The more frequent sampling of processing-competent non-s1 states by the YA6 mutant in the absence of substrate is also consistent with its higher basal ATPase rate. Interestingly, the YA1-mutant proteasome did not show such a shift in the conformational distribution (Supp. Fig. 2.3-2), and its increased basal ATP hydrolysis may be caused by an effect distinct from that observed for the YA6 mutant. Based on our data, Rpt6 appears particularly important for sensing the absence or presence of a substrate polypeptide in the central channel and controlling the proteasome’s conformational transitions. Structures of various other AAA+ protein translocases in the presence of substrate revealed a network of interactions between conserved pore-1 loop residues, in which a positively charged Lys or Arg preceding the grip-conferring tyrosine interacts with a negatively charged Glu or Asp in the pore-1 loop of the neighboring ATPase subunit in the ring ^41^. Furthermore, pore-1-loop interactions were suggested to gate the central channel prior to substrate insertion, and it is possible that mutation of Rpt6’s tyrosine disrupts these interactions and partially decouples s1-to-non-s1 conformational changes from substrate engagement.

**Fig. 3.**
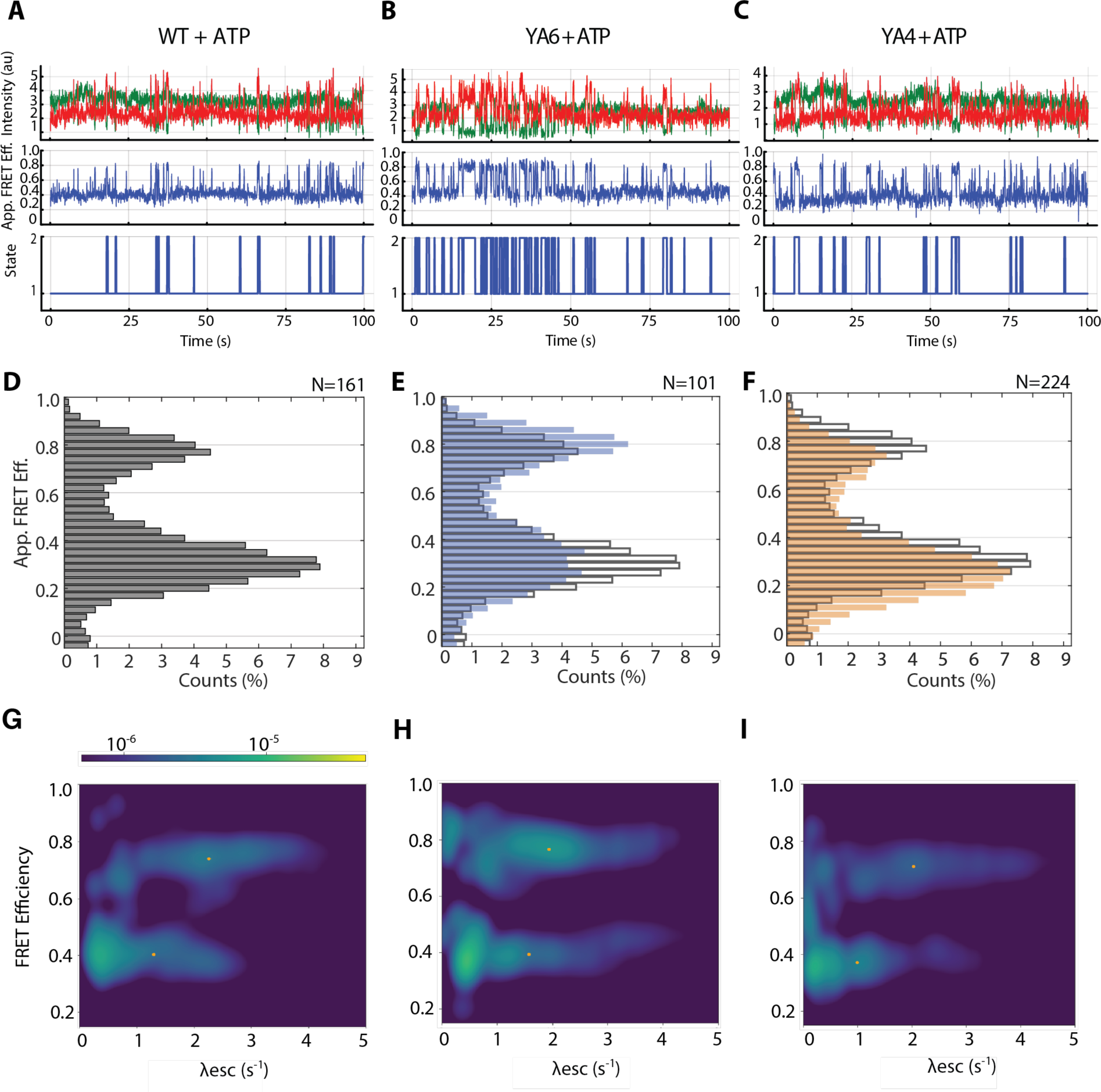
Differential effects of Rpt4 and Rpt6 pore-1 loop mutations on the proteasome conformational dynamics. **A, B, C)** Example traces for wild-type (A), YA6 (B), and YA4 proteasomes (C) in ATP, with the donor (green) and acceptor (red) fluorescence intensities shown in the top panels, the apparent FRET efficiencies in the middle panels, and estimated most-probable state trajectories in the bottom panels. **D)** Apparent FRET-efficiency distributions for wild-type proteasomes in ATP. **E, F)** Apparent FRET efficiency distributions for YA6 (blue bars) and YA4 (orange bars) mutant proteasomes overlayed with the distribution of wild-type proteasome (grey outlined bars). **G, H, I)** Bivariate probability distributions for the escape rates (λesc) of wild-type, YA6, and YA4 mutant 26S proteasomes in ATP, respectively. The logarithmic scale color map (see top bar) describes probability regions that indicate the likelihood of proteasomes to assume a specific FRET efficiency based on all sampled states.

To understand the effect of pore-1 loop mutations on substrate processing, we measured the single molecule conformational dynamics of the proteasome in the presence of a model substrate, ubiquitinated titin I27^V15P^. We previously found that upon substrate engagement the proteasome stably switches to processing-competent, high-FRET states that are maintained until substrate unfolding and threading through the ATPase motor has been completed ^38^. With the new instrumental setup, we first confirmed these findings by observing high-FRET phases, i.e. substrate-processing dwells, and apparent FRET efficiency distributions for the wild-type proteasome in the presence of ubiquitinated titin I27^V15P^ substrate that were comparable to the previously reported values (Fig. 4A,D; Supp. Fig. 4-1, 4-2, 4-3). Traces for the YA4 mutant exhibited much shorter dwells, which were caused by transient low-FRET-state excursions interrupting the high-FRET degradation phase (Fig. 4C, Supp. Fig. 4-1C). Our previous investigations of conformational transitions revealed that the wild-type proteasome also shows brief low-FRET excursions, with a frequency that positively correlates with the substrate’s thermodynamic stability ^38^. These transitions may thus represent slippage events and temporary returns to the s1 state after unsuccessful substrate-unfolding attempts by the motor.

**Fig. 4.**
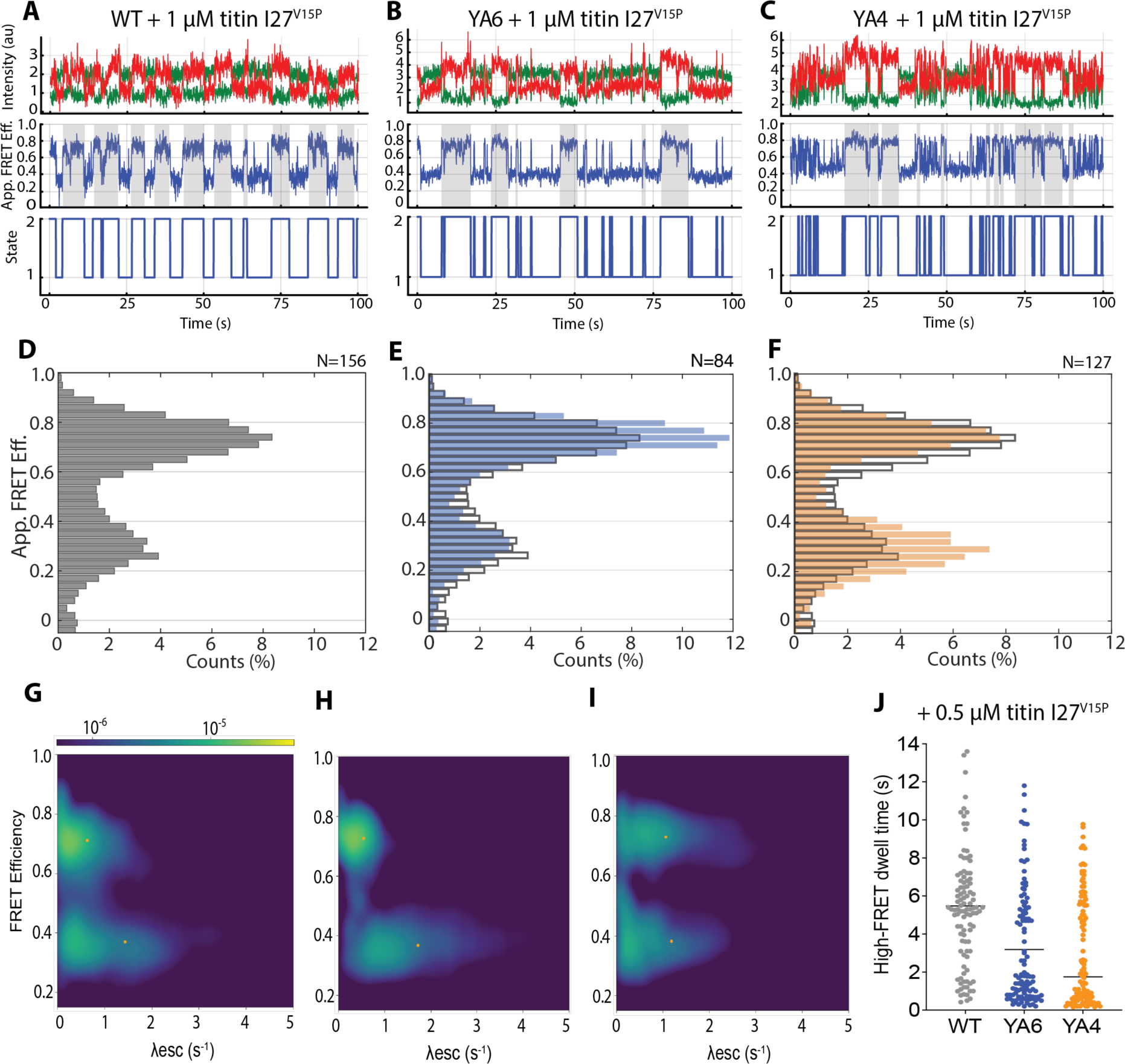
The YA4 and YA6 mutations lead to frequent transitions out of the processing-competent states during substrate degradation. **A, B, C)** Example traces for wild-type (A), YA6 (B), and YA4 mutant 26S proteasomes (C) in the presence of 1 μM ubiquitinated titin I27^V15P^ substrate, with the donor (green) and acceptor (red) fluorescence intensities shown in the top panels, the apparent FRET efficiencies in the middle panels, and estimated most-probable state trajectories in the bottom panels. High-FRET dwells during substrate processing are highlighted by grey shading. **D)** Apparent FRET efficiency distributions for wild-type proteasomes 1 μM ubiquitinated titin I27^V15P^ substrate. **E, F)** Apparent FRET efficiency distribution for YA6 proteasomes (blue bars) and YA4 proteasomes (orange bars) in the presence of 1 μM ubiquitinated titin I27^V15P^ substrate, compared to the distribution for wild-type proteasomes (grey outlined bars). **G, H, I)** Bivariate probability distributions for the escape rates (λesc) of wild-type, YA6, and YA4 proteasomes, respectively, in the presence of 1 μM ubiquitinated titin I27^V15P^ substrate. The logarithmic scale color map (see top bar) describes probability regions that indicate the likelihood of proteasomes to assume a specific FRET efficiency based on all sampled states. **J)** Lengths of continuous high-FRET phases during the degradation of ubiquitinated titin I27^V15P^ substrate by wild-type, YA6, and YA4-mutant proteasomes. The substrate concentration for these analyses was 0.5 μM for larger spacing and thus better discrimination of individual substrate-processing events.

The YA6 mutant in the presence of titin I27^V15P^ substrate resembled the wild-type proteasome in its overall FRET-efficiency distribution (Fig. 4D, 4E), and we used the above-described HMM-MCMC methods to determine the kinetics of conformational transitions during degradation of 1 μM titin I27^V15P^ substrate. In these analyses, YA6 again resembled wild-type proteasome in the mean rates for escape from the low-FRET and high-FRET states (1.72 and 0.54 s^-1^ for YA6 versus 1.43 and 0.63 s^-1^ for WT; Fig. 4G, 4H, Supp. Table 1). Similar to YA6, the YA1, YA2, YA3, and YA5-mutant proteasomes showed wild-type-like FRET-efficiency distributions (Supp. Fig.4-2). In contrast, the YA4-mutant exhibited a pronounced shift in its FRET-efficiency distribution toward the low-FRET s1 state (Fig. 4F) and a correspondingly higher rate of 1.05 s^-1^ for the escape from the high-FRET states (Fig. 4I), consistent with the frequent excursions to the low-FRET state that we observed in individual traces during substrate processing.

The FRET-efficiency traces for proteasomes in the presence of substrate contain phases of active degradation as well as phases between degradation events during which the proteasome is not occupied with substrate. To more specifically analyze how the YA mutations affect the conformational transitions during active degradation, we therefore manually scored the high-FRET phases of proteasomes in the presence of 0.5 μM ubiquitinated titin I27^V15P^ substrate. The wild-type proteasome showed an average length of 1^WT^ = 5.5 ± 0.3 s for the persistence of high-FRET, processing-competent states, whereas YA4 and YA6-mutant proteasomes had shorter high-FRET phases, with 1^YA4^ = 3.2 ± 0.3 s and 1^YA6^ = 3.1 ± 0.3 s (Fig. 4J, Supp. Table 2). We therefore hypothesize that the absence of the pore-1 loop tyrosine in Rpt4 and Rpt6 reduces the motor grip, causes slippage during substrate-unfolding attempts, and leads to recurring transitions to the s1 state, potentially accompanied by disengagement from the substrate in the central channel.

In summary, our data revealed that the pore-1 loop mutation in Rpt6 influences the proteasome conformational dynamics in the absence of substrate, and mutation of both Rpt6’s and Rpt4’s pore-1 loop affects the stable maintenance of the processing-competent states during substrate unfolding, potentially due to slippage.

### Pore-1 loops of Rpt4 and Rpt6 play distinct roles in substrate processing

While YA mutations in Rpt1, Rpt2, Rpt3, and Rpt5 only minimally affected substrate degradation, YA mutations in Rpt4 and Rpt6 led to major defects. We therefore focused on these two subunits and employed a previously developed FRET-based substrate-procession assay to elucidate how their pore-1 loops contribute to the individual steps of degradation ^17,38^. The FRET-based assay relies on fluorophores attached to the substrate’s flexible initiation region and to an AzF substituted for Ile191 in the linker between the N-domain and ATPase domain of Rpt1, allowing to directly monitor the progression of a substrate polypeptide through the central channel (Fig. 5A). The dyes were placed such that initial substrate binding to the proteasome leads to an intermediate FRET efficiency, which increases to a maximum value after substrate-tail insertion, engagement by the ATPase motor, and the onset of translocation. The high FRET efficiency then persists during deubiquitination, before declining as the substrate is translocated further through the ATPase channel and into the 20S CP ^38^.

**Fig. 5.**
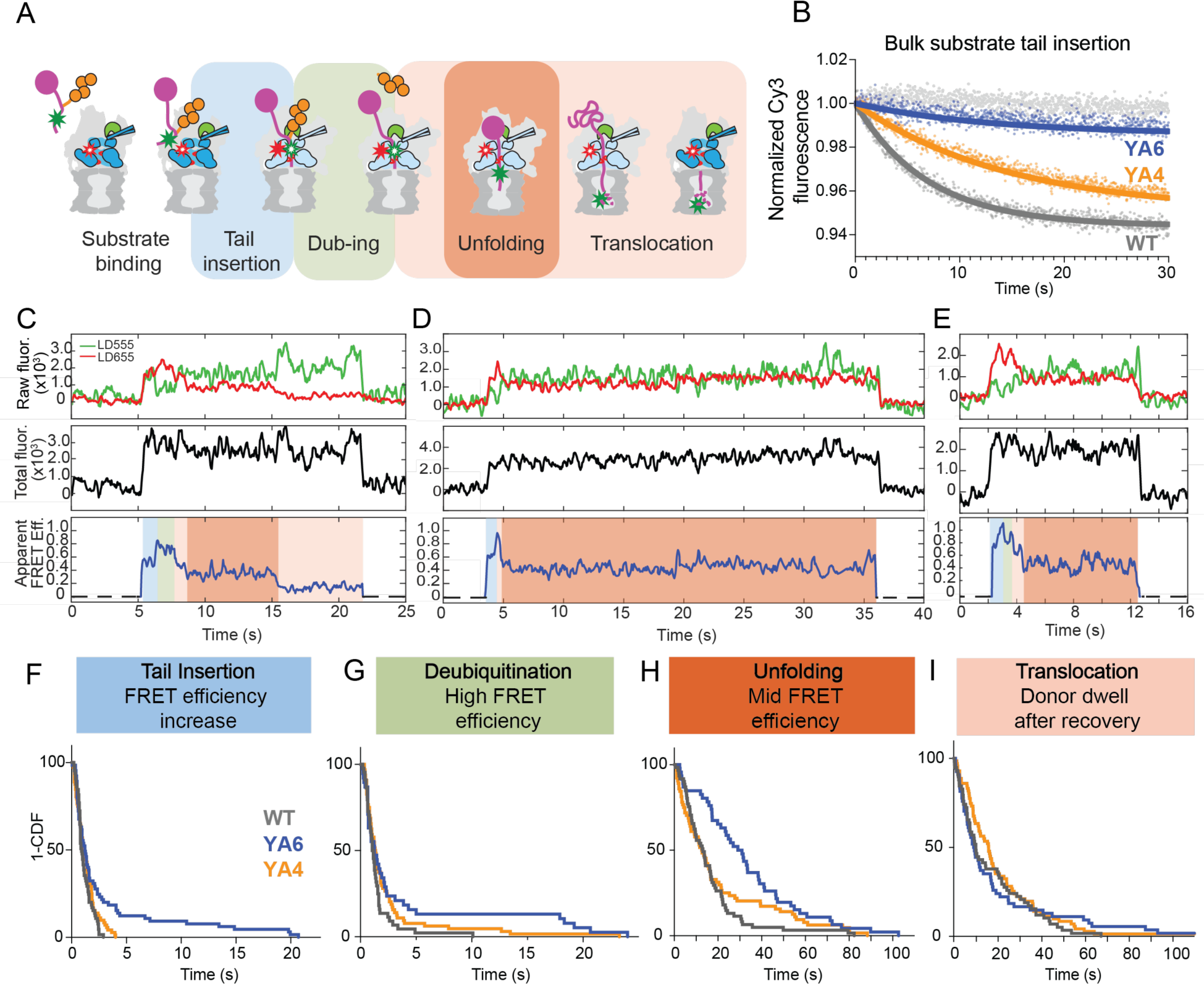
Direct observation of substrate processing by pore-1 loop mutant proteasomes using single-molecule FRET. **A)** Schematic for the FRET-based substrate-processing assay, monitoring the individual steps of substrate binding and tail insertion (blue shading), deubiquitination (green shading), unfolding (dark orange shading), and translocation (light orange shading). **B)** Representative traces and single exponential fits for the bulk substrate-processing assay with Cy5-acceptor-labeled substrate and Cy3-donor-labeled proteasomes whose Rpn11 deubiquitinase was inhibited to stall substrate processing after tail insertion and engagement. Shown are the changes in Cy3 donor fluorescence after stopped-flow mixing with ubiquitinated titin I27^V15P^ substrate for wild-type (dark grey), YA6 (blue), and YA4 proteasomes (orange). The data were normalized to initial fluorescence values. A control with ubiquitinated, Cy5-labeled titin I27^V15P^ substrate and wild-type Cy3-labeled base subcomplex alone, in the absence of lid and core peptidase, is depicted in light grey. **C)** Representative single-molecule fluorescence traces for the degradation of LD555 donor-labeled ubiquitinated titin I27 substrate by LD655 acceptor-labeled 26S proteasomes. The top panel shows fluorescence intensities for the FRET donor (green) and acceptor (red), the middle panel shows the calculated total fluorescence intensity, and the bottom panel illustrates the calculated apparent FRET efficiency. Substrate degradation events showed a constant total fluorescence, indicating the presence of a single substrate per proteasome. Traces were filtered for visualization using Matlab R2019b. **D, E)** Example traces for titin I27 substrate processing by YA6 (D) and YA4 mutant proteasomes (E) show longer dwells for the unfolding phase. **F)** Cumulative distribution function (1-CDF) or survival plot analysis for the tail insertion and engagement step of wild-type proteasome (grey, N = 65), the YA6 mutant (blue, N = 66), and the YA4 mutant (orange, N = 72). **G)** Survival plot analysis for the deubiquitination step of wild-type proteasome (grey, N = 44), the YA6 mutant (blue, N = 39), and the YA4 mutant (orange, N = 64). **H)** Survival plot analysis for the unfolding step of wild-type proteasome (grey, N = 61), the YA6 mutant (blue, N = 48), and the YA4 mutant (orange, N = 64). **I)** Survival plot analysis for the translocation step after the unfolding dwell for wild-type proteasome (grey, N = 58), the YA6 mutant (blue, N = 54), and the YA4 mutant (orange, N = 71).

First, we used this assay in stopped-flow mixing experiments with sulfo-Cy3 donor-dye labeled proteasomes and sulfo-Cy5 acceptor-dye labeled ubiquitinated titin I27^V15P^ substrate to monitor the kinetics of substrate-tail insertion and engagement in bulk ^17^. For these ensemble measurements, reconstituted proteasomes were incubated with the Rpn11 inhibitor 1,10-phenanthroline (oPA) to stall substrate processing at the high-FRET deubiquitination phase right after engagement, leading to a single-turnover scenario in which the combined first-order time constant for substrate binding, tail insertion, and engagement could be determined by fitting the increase in FRET-efficiency (Fig. 5B, Supp. Fig. 5-1). This time constant was 1_ins_^WT^ = 6.8 ± 0.1 s for wild-type proteasomes, whereas YA4- and especially YA6-mutant proteasomes showed significant delays, with time constants of 1^insYA4^ = 14.3 ± 0.8 s and 1_ins_^YA6^ = 18.0 ± 2.5 s, respectively. The almost 3-fold slower substrate insertion into the YA6 mutant can be explained with its shift of the conformational equilibrium and frequent spontaneous sampling of the processing-competent non-1 states, in which the Rpn11 deubiquitinase obstructs the entrance to the motor channel and thus inhibits substrate insertion. Interestingly, the YA6 mutant not only exhibited slower kinetics, but also a 4.5-fold lower amplitude than the wild-type proteasome in the FRET-signal change, suggesting that substrates either do not fully enter the ATPase ring or fail to stay stably inserted. Lack of the Rpt6 pore-1 loop tyrosine thus not only delays substrate insertion, but also appears to compromise substrate capture, likely due to defects in motor engagement of the substrate’s initiation region.

To gain more detailed insights into these engagement problems and discern potential defects in mechanical unfolding, we switched to the thermodynamically more stable wild-type titin I27 substrate. After confirming robust degradation of this substrate by wild-type and YA-mutant proteasomes in bulk (Supp. Fig. 5-2), we used the FRET-based substrate-processing assay in our upgraded single-molecule TIRF microscopy setup for further investigation. In this single-molecule assay, the substrate was LD555-donor labeled, and immobilized proteasomes carried a LD655 acceptor. Fluorescence traces for this experimental setup show an initial intermediate FRET-efficiency value of ∼ 0.5 upon binding of the ubiquitinated substrate to a proteasomal receptor, followed by a FRET increase to its maximum value of 0.8 - 1.0 when the substrate tail inserts into the ATPase motor, and a short high-FRET dwell during Rpn11-mediated deubiquitination (Fig. 5C). Further substrate translocation then leads to a decrease in the apparent FRET-efficiency signal to a persistent value of 0.35 - 0.45 that we identified as a pre-unfolding dwell, which was much harder to observe previously when using the more labile and readily unfolded titin I27^V15P^ substrate. Successful unfolding and continuous translocation subsequently results in a FRET-efficiency decay to ∼ 0.1 – 0.2, before proteolytic cleavage and release of the donor-labeled fragment from the internal chamber leads to loss of the donor signal.

We analyzed the FRET-efficiency traces for wild-type and YA-mutant proteasomes during titin I27 degradation by scoring the processing steps represented by the individual FRET phases and plotting their cumulative distribution functions (1-CDF or survival plots, Fig. 5 F-I, Supp. Fig. 5-3). Fitting the survival plots for the tail insertion phase to single-exponential functions revealed that the majority of particles for all YA mutants showed wild-type-like kinetics, with time constants ranging from 0.8 to 1.6 s (Supp. Table 3). Only a small fraction of ∼ 10 % for the YA6-mutant proteasome took on the substrate with delayed kinetics (Fig. 5F). Successful tail insertion thus occurs largely with similar kinetics even for the engagement-defective YA6 mutant, suggesting that the defects we observed in bulk measurements originate from failed insertion attempts that are not considered in the survival plots (see below). The time constants for the subsequent deubiquitination phase were again similar for all YA mutants and wild type, ranging between 0.8 s and 1.5 s (Supp. Table 4). Interestingly, substrate unfolding also occurred at wild-type-like rates for all YA mutant except YA6, which spent 3-fold more time on unraveling titin I27 (1 ^YA6^ = 50.8 s versus 1 ^WT^ = 16.3 s; Fig. 5D,H, Supp. Table 5). This observation for the YA6 proteasome is remarkable, as we detected no obvious unfolding defect for this mutant during degradation of the less stable titin I27^V15P^ in our single-molecule conformational dynamics and substrate-processing assays (Fig. 4B,E,H; Supp. Fig.5-4, Supp. Table 7). Manifestation of YA6’s unfolding defects thus appears to depend on the substrate’s thermodynamic stability and possibly the motor’s behavior during unfolding attempts. Also surprising is that the YA4 mutant showed no extended unfolding phase, suggesting that its degradation defects are not necessarily caused by slower unfolding, but possibly lower processivity and premature release (see below). The subsequent substrate-translocation phase, identified as the donor dwell after successful substrate unfolding and FRET-efficiency decay to ∼ 0.1 – 0.2, was similar for wild-type and most of the pore-1 loop-mutant proteasomes, with a slightly longer time constant for YA4 and faster translocation for the YA1 mutant (Fig. 5I, Supp. Fig.5-3, Supp. Table 6).

### Efficient substrate capture depends on both Rpt6 and Rpt4

Since the YA6-mutant proteasome exhibited substrate-tail insertion and engagement defects in our ensemble measurements that were not as obvious when monitoring successful processing events by single-molecule FRET, we analyzed the success rate of this mutant in capturing the titin I27 substrate compared to wild-type and other YA-mutant proteasomes. The capture success was quantified based on at least 250 events of substrate interactions, for which we calculated the ratio of successful substrate engagement and processing relative to the total number of substrate-binding events, as previously described ^38^. These calculations yielded a capture-success rate of 51.1 ± 5.1 % for the wild-type proteasome and similar values for the YA1, YA2, YA3, and YA5 mutants, indicating no defects in substrate engagement for these variants (Fig. 6A, Supp. Fig. 6-1A, Supp. Table 8). However, the YA6-mutant proteasome showed a capture success of only 26.1 ± 5.3 %, and substrate capture of the YA4 mutant was with 33.9 ± 4.5 % also considerably diminished (Fig. 6A). The almost 2-fold reduced ability of the YA6 mutant to successfully engage a substrate before it dissociates from a ubiquitin receptor is consistent with the observed shift in this mutant’s conformational equilibrium and the ∼ 2-fold reduced abundance of the engagement-competent s1 state (Fig. 3E). In contrast, the YA4 mutant did not show an equivalent shift in the conformational equilibrium. Insights into potential differences between the underlying reasons for the substrate-capture problems of YA6 and YA4-mutant proteasomes was provided by analyzing the residence time of substrates on the proteasome during unsuccessful capture attempts. These interaction were short for the YA6-mutant with an average time constant of τ = 0.4 s, whereas substrates spent ∼ 2-fold more time on the YA4-mutant proteasome before dissociating (τ = 0.8 s; Supp. Fig.6-1C). While the lack of an intact Rpt6 pore-1 loop seems to cause early defects by conformationally interfering with substrate-tail insertion, mutation of Rpt4’s pore-1 loop may prevent a stable commitment to substrate degradation at a slightly later stage, for instance due to failed power strokes that drive substrate translocation after engagement. As expected, measurements with the less stable titin I27^V15P^ substrate revealed a similar picture, with a capture success of 51.2 ± 4.6 % for wild-type proteasome and 34.9 ± 0.2 % and 36.6 ± 2.3 % for the YA4 and YA6 mutants, respectively (Supp. Fig. 6-1B, Supp. Table 8).

**Fig. 6.**
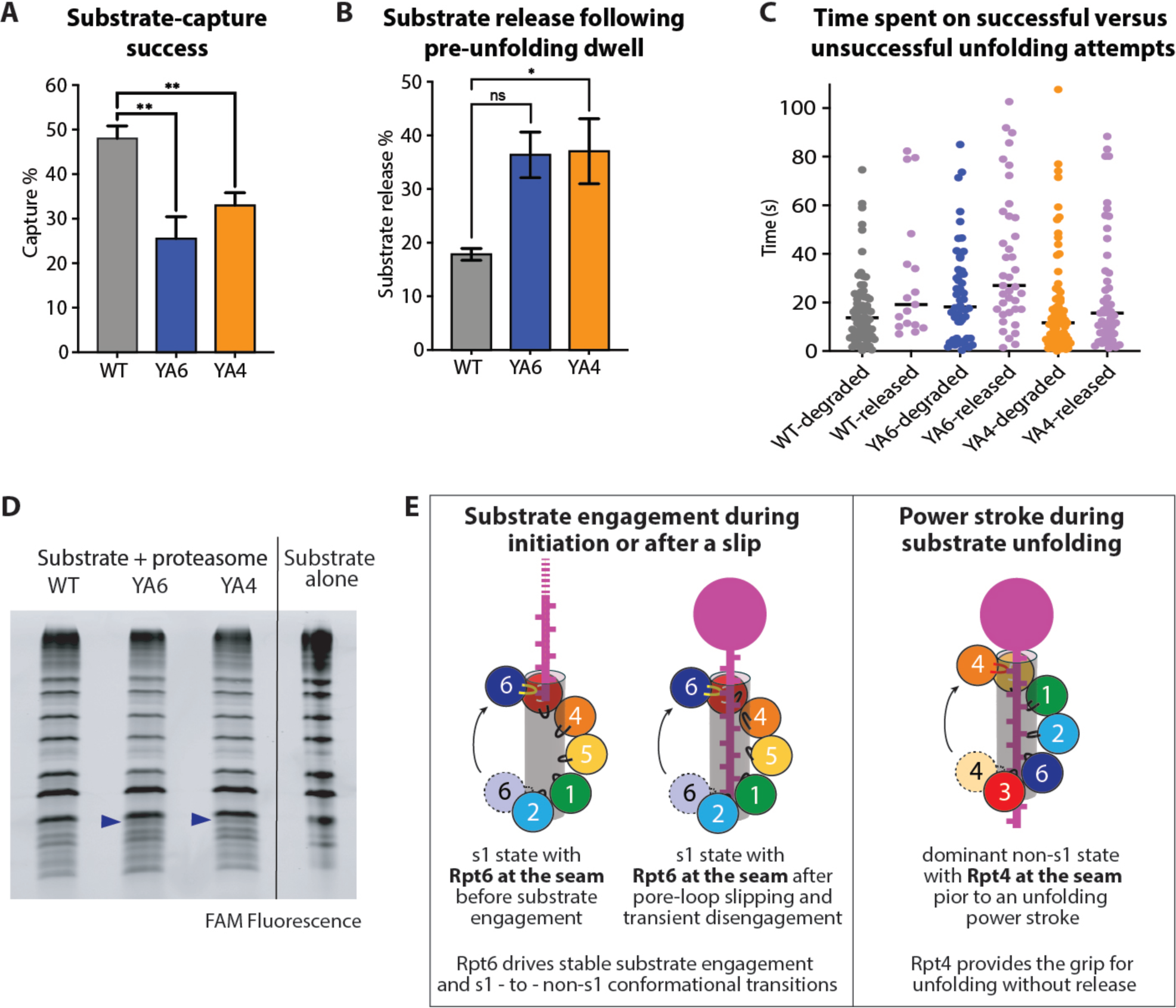
The pore-1 loops of Rpt6 and Rpt4 are critical for efficient substrate capture and unfolding without release. **A)** Success rates for the capture of titin I27 substrate by wild-type (grey), YA6 (blue), and YA4 mutant proteasome (orange). N = 2 technical replicates for all proteasome variants, with each contributing at least 250 capture events. Statistical significance was calculated using a one-way ANOVA test: **p<0.01; *p<0.05; ns, p>0.05. **B)** Percentage of titin I27 release during the unfolding dwell as determined by single-molecule FRET for wild-type (grey), YA6 (blue), and YA4 mutant proteasome (orange). N ≥ 2 technical replicates with each contributing to at least 102 events. **C)** Distributions of time spent by wild-type proteasome (grey), the YA6 mutant (blue), and the YA4 mutant (orange) on successful substrate unfolding attempts versus unsuccessful attempts that were terminated by substrate release during the unfolding dwell (magenta), as determined by single-molecule FRET (N = 93, 91, and 91 total events for wild-type, YA6, and YA4 mutant proteasomes, respectively). **D)** SDS-PAGE analysis of the 60-min end point samples from the multiple-turnover degradation of ubiquitinated FAM-labeled titin I27 substrate by wild-type, YA6, and YA4-mutant proteasomes. The accumulation of deubiquitinated and truncated products (indicated by arrowheads) was visualized by FAM fluorescence. **E)** Model for the critical roles of Rpt6 and Rpt4 in substrate engagement and processing. Left and middle: Rpt6 is always at the seam position in the s1-state spiral staircase of Rpts and therefore the first subunit to either engage a substrate during degradation initiation or re-engage a substrate during processing after slipping caused transient pore-loop disengagement and a brief return to the s1 state. Right: The strongly dominant non-s1 state for the substrate-degrading proteasome has Rpt4 in the seam position of the spiral staircase. Rpt4 is therefore the first subunit to grab the substrate at the top of the staircase during an unfolding power stroke, which may be followed by rapid firing of the other Rpt subunits in a hand-over-hand mechanism.

### Rpt4 and Rpt6 ensure robust substrate unfolding without release

Interestingly, many FRET traces for titin I27 degradation by the YA4- and YA6-mutant proteasomes did not show a final translocation phase, but ended abruptly with a loss of the donor signal while the proteasomes resided in the pre-unfolding phase (Fig. 5D,E), suggesting that unsuccessful unfolding attempts led to slippage and terminal substrate release. The hypothesis that motor slippage could lead to substrate backsliding in the central channel is supported by interesting FRET-efficiency patterns that we observed for several YA-mutants and even the wild-type proteasome during titin I27 degradation, where multiple transient high-FRET peaks interrupted the persistent 0.35 - 0.45 FRET efficiency phase in the pre-unfolding dwells (Supp. Fig. 5-5). This appear to be the first direct observation of substrate slippage, where loss of grip by the pore-1 loops may cause the unstructured substrate tail to backslide, bringing the attached donor dye again into closer proximity to the acceptor dye above the ATPase ring.

To further investigate the contributions of individual pore-1 loops to unfolding and motor grip on the substrate, we analyzed ∼ 100 traces for wild-type and YA-mutant proteasomes, and determined their frequency of terminal substrate release after unsuccessful unfolding attempts, i.e. events where substrate processing ended abruptly with a loss of donor fluorescence during the unfolding dwell (Fig. 6B, Supp. Fig. 6-2A). In control measurements with immobilized LD555-labeled proteasomes we determined the fluorescence lifetime of the LD555 donor dye as 1_bleach_^LD555^ = 127 s (Supp. Fig. 6-2B), which is much longer than most of the pre-unfolding dwell times and thus confirms that the abrupt termination of substrate-processing events was indeed due to substrate release rather than photobleaching of the donor dye. Wild-type proteasome released the titin I27 substrate in only 16.4 ± 0.2 % of degradation attempts, and the YA1, YA2, YA3, and YA5 mutants were equally efficient in substrate unfolding (Supp. Fig.6-1C, Supp. Table 9). Importantly, however, YA4 and YA6-mutant proteasomes showed more than 2-fold higher release frequencies, letting go of 38.0 ± 7.3 % and 35.6 ± 3.1 % of their substrates (Fig. 6B).

Furthermore, we analyzed for how long different proteasome variants tried to unfold the titin I27 substrate before either succeeding or releasing it. We found that it takes wild-type, YA6, and YA4-mutant proteasomes overall similar amounts of time for successful titin I27 unfolding (Fig. 6C). While the YA4 mutant showed a similar time distribution for unsuccessful attempts, with many releases occurring already shortly after entering the unfolding phase, the YA6-mutant proteasome spent ∼ 2 times longer on unsuccessful unfolding attempts before release (Fig. 6C). These ultimately unsuccessful attempts are therefore responsible for the apparently longer unfolding times of the YA6 mutant (Fig. 5H). Overall, we can conclude that the substrate-unfolding defects of YA6- and YA4-mutant proteasomes primarily originate from their significantly increased release frequency. The YA6 mutant thereby spends more time on unfolding attempts, whereas the YA4 mutant more readily releases the substrate, suggesting an overall lower grip during unfolding power strokes. Unsuccessful unfolding and release of the wild-type titin I27 substrate by YA4 and YA6-mutant proteasomes is expected to cause some accumulation of deubiquitinated substrate species with slightly truncated initiation regions. Indeed, we were able to confirm the increased presence of these products for the YA4 and YA6 mutants by SDS-PAGE analyses of the end points from multiple-turnover degradation reactions (Fig. 6D).

## Discussion

Here, we uncovered that particular pore-1 loops in the proteasomal heterohexameric AAA+ ATPase motor play specific roles for substrate engagement, unfolding, and the conformational response of the 19S RP. By measuring the proteasome conformational dynamics and directly observing the individual steps of substrate processing, we found that the pore-1 loop of Rpt6 is especially critical for substrate capture and stable engagement, as well as controlling the transitions between engagement-competent and processing-competent proteasome states. Removal of Rpt6’s pore-1 loop tyrosine shifts the conformational equilibrium in the absence of substrate toward processing-competent non-s1 states. Due to the obstruction of the channel entrance by the centrally localized Rpn11 deubiquitinase in non-s1 states, this conformational shift interferes with substrate insertion and lowers the success rate of capturing ubiquitinated substrates for degradation ^17,38^. These findings indicate that Rpt6’s pore-1 loop stabilizes the engagement-competent s1 state in the absence of substrate and/or facilitates the transition to non-s1 states upon motor engagement of a substrate polypeptide, potentially through interactions of its conserved tyrosine residue with the pore-1 loops of neighboring Rpt subunits. The importance of Rpt6’s pore-1 loop for stable substrate engagement can be explained with its position in the spiral-staircase arrangement of Rpt subunits in the engagement-competent s1 state. Only a single staircase register has been observed for the substrate-free, engagement-competent proteasome, with Rpt3 at the top, Rpt2 at the bottom, and Rpt6 in an intermediate “seam” position of the staircase ^21–29,31–37^. Rpt6 is therefore expected to always be the first subunit to move up to the top of the staircase, engage the inserted substrate polypeptide, and hence drive the commitment step, in which the 19S RP transitions from the s1 to non-s1 states and the system switches from passive substrate diffusion to active, ATP-hydrolysis-driven translocation (Fig. 6E). Insufficient steric interactions during this first step of engaging a newly inserted substrate would explain the substrate capture and engagement defects that we detected for the YA6 mutant in our single-molecule and bulk measurements.

The importance of Rpt6’s pore-1 loop for substrate engagement also explains the unfolding defects observed for the YA6 mutant. Our previous single-molecule studies revealed that during unfolding of tough substrates, the proteasome transiently switches from the processing non-s1 states to the s1 state, possibly due to pore-loop slippage and brief disengagement from the substrate polypeptide in the central channel ^38^. It is conceivable that subsequent transitioning back to non-s1 states for continued unfolding attempts and translocation involves pore-loop re-engagement, in which Rpt6 as the seam subunit in the s1-state spiral plays again a particularly important role (Fig. 6E). Failure to rapidly re-engage the substrate would lead to delays in unfolding and to substrate escape from the central channel, as we observed for the YA6 mutant.

While the YA6-mutant proteasome appears to have problems re-engaging a substrate and returning to non-s1 processing states after a slip, the YA4 mutation seems to cause more frequent slips and s1-state transitions that interrupt substrate unfolding, suggesting a reduced grip during the power stroke. This hypothesis is well supported by findings from several cryo-EM structural studies of substrate-engaged human and yeast 26S proteasomes, which consistently showed the vast majority of particles (∼ 60 – 80 %, Supp. Table 10) in spiral staircase orientations with Rpt4 at either the bottom or the seam position and occupied with ADP or no nucleotide ^31,37,42^. This spiral state is therefore 4-5-fold more prevalent than expected if each register was equally likely to be observed during a hand-over-hand translocation mechanism. For the cryo-EM structure determinations, proteasomal substrate translocation was transiently stalled either on an uncleaved ubiquitin chain, a stably folded substrate domain, or upon addition of the non-hydrolysable ATP analog ATPγS, and it is possible that the dominant staircase orientation with Rpt4 at the bottom or the seam positions represents a pre-stroke state when the motor is dealing with a tough barrier. In this staircase register, Rpt4 would be the next subunit to move up, contact the substrate polypeptide at the top of the Rpt spiral, and drive the subsequent power stroke (Fig. 6E). Lack of Rpt4’s pore-1 loop tyrosine may therefore reduce the grip during unfolding attempts, cause more frequent motor slippage, and substrate release, consistent with our experimental data for the YA4 mutant.

For distinct reasons, YA6 and YA4-mutant proteasomes both reduce the substrate-capture success by about 2-fold and increase substrate release during unfolding attempts by another factor of 2, leading to overall major defects in substrate processing. In contrast, removing the pore-1 tyrosines from any of the other four Rpt subunits has only minor effects on degradation. Our results therefore indicate asymmetric mechanisms for the proteasomal AAA+ motor in which subunits at the bottom or seam positions of particular spiral-staircase arrangements in the ATPase ring play critical roles for substrate capture and robust unfolding without release. Based on these biochemical findings in combination with previous cryo-EM structural data ^31,37,42^, we propose an adaptation to the current hand-over-hand translocation mechanism. Rather than uniformly progressing through the ATP-hydrolysis cycle and the different registers of the spiral staircase, the Rpt subunits may adopt a particular staircase orientation when encountering an unfolding barrier, with Rpt4 at the bottom or seam position. Coordinated, sequential ATP hydrolysis events and conformational changes may then rapidly progress around the ring until Rpt4 is in the bottom or seam position again. Such firing in bursts would explain the strongly skewed distribution of spiral staircase states observed in the cryo-EM structural studies of the 26S proteasome. This mechanism would also be consistent with the larger, 10 – 40 Å increments of translocated substrate polypeptide that were measured in optical tweezing experiments during substrate degradation by the related bacterial AAA+ protease ClpXP and may originate from the coordinated firing of several subunits in rapid bursts ^43,44^. Although future, more detailed and time-resolved biophysical studies will be required to confirm that the 26S proteasome and potentially other AAA+ motors use such bursts in ATP hydrolysis, our results presented here clearly reveal the importance asymmetric mechanisms for efficient substrate engagement and unfolding by the proteasomal ATPase motor and thus provide an important advance compared to structural snapshots of the hand-over-hand mechanism.

## Supplemental Figures

**Supp. Fig.1-1:**
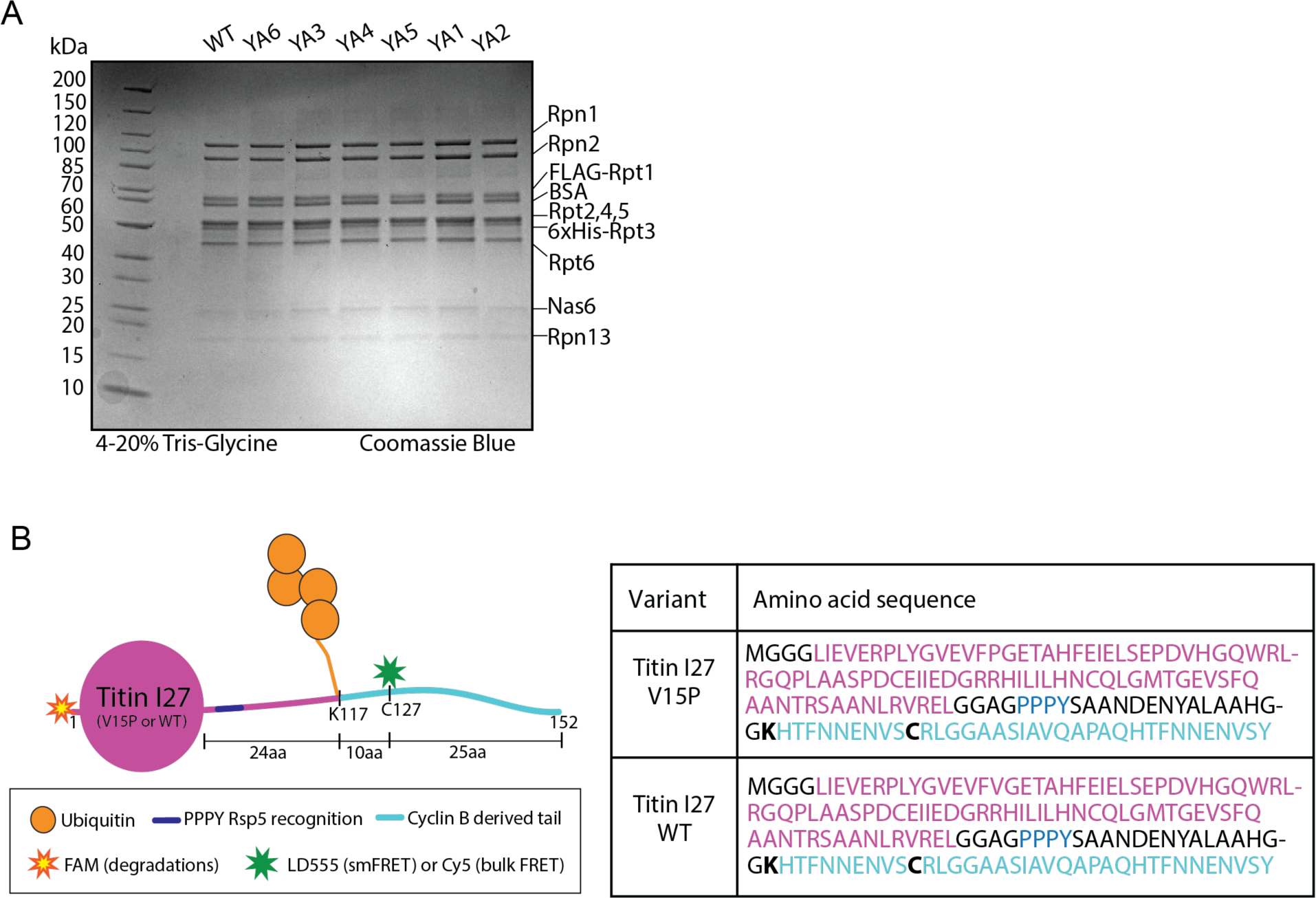
**A)** 4-20% Tris-Glycine SDS-PAGE of purified base subcomplexes (∼3.2 μg), visualized by Coomassie blue stain. **B)** Titin I27 substrate used throughout this study, either without (wild type) or with a V15P destabilizing mutation as indicated in each assay. All lysine residues were removed from the folded domain, and a single lysine for ubiquitin attachment was introduced at position 117 within a C-terminally fused Cyclin B-derived unstructured tail (cyan). An Rsp5 E3 ligase recognition site (dark blue) was added to facilitate polyubiquitination (orange circles). In addition, the substrate was labeled either at the N-terminus through a Sortase A reaction with a FAM-LPETGG peptide, used to track substrate degradation, or at an engineered Cys127 with maleimide-linked SulfoCy5 or LD555 to follow substrate processing by FRET in bulk or at the single-molecule level, respectively. Sequences for the wild-type and V15P titin substrates are shown on the right.

**Supp. Fig.1-2:**
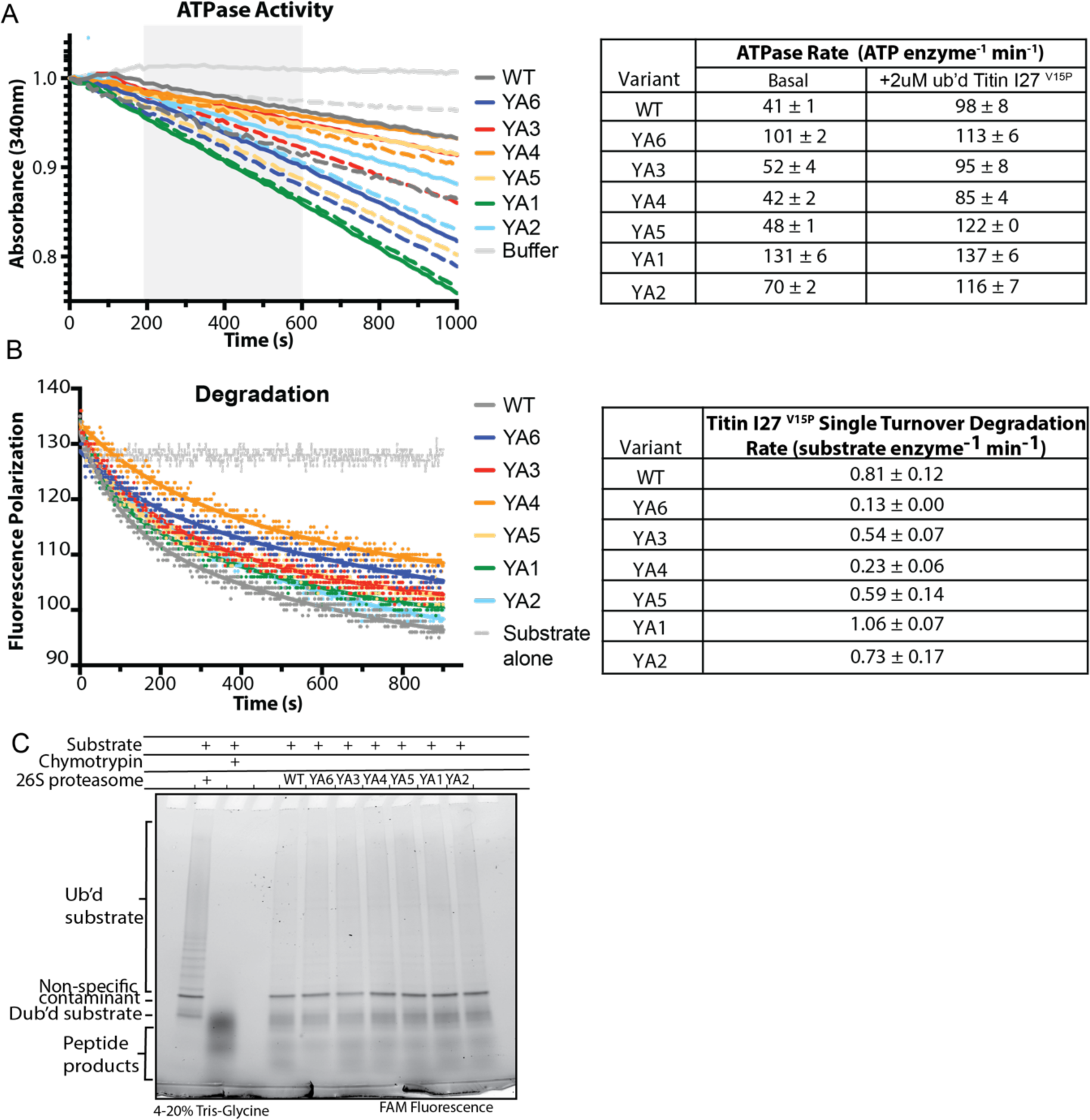
**A)** Representative traces for the ATP-hydrolysis measurement of *in vitro*-reconstituted proteasomes in the absence (dashed lines) or presence (solid lines) of 2 μM ubiquitinated FAM-titin I27^V15P^ in a NADH-coupled assay. The shaded area was used for linear regressions of the observed absorbance decays, with the resulting ATPase rates shown in the table on the right. **B)** Representative traces for the single-turnover degradation of the ubiquitinated FAM-Titin I27^V15P^ substrate by *in vitro*-reconstituted proteasomes, monitored by the decrease in fluorescence polarization. Data were fit to a double exponential decay (solid line) using GraphPad Prism to obtain degradation rates shown in the table on the right. **C)** SDS-PAGE analysis of end- point samples for the single-turnover degradation of the ubiquitinated FAM-Titin I27^V15P^ substrate visualized using the FAM fluorescence emission channel.

**Supp. Fig.2:**
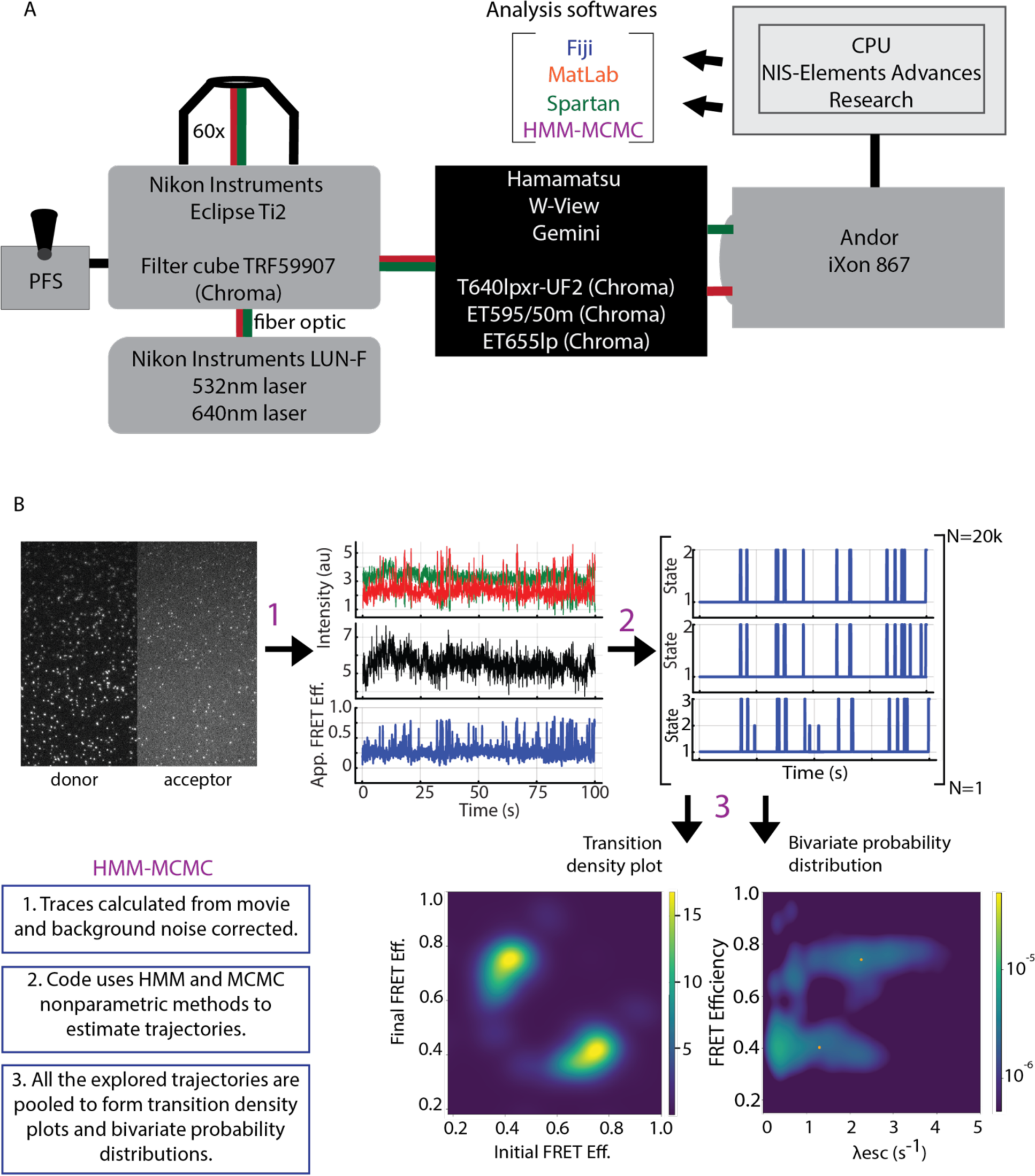
**A)** TIRF-microscope setup using the Nikon Instruments Eclipse Ti2, LUNF laser box, and Perfect Focus system. **B)** Hidden Markov Model - Markov Chain Monte Carlo algorithm and workflow for the analyses of the proteasome conformational dynamics.

**Supp. Fig. 3-1.**
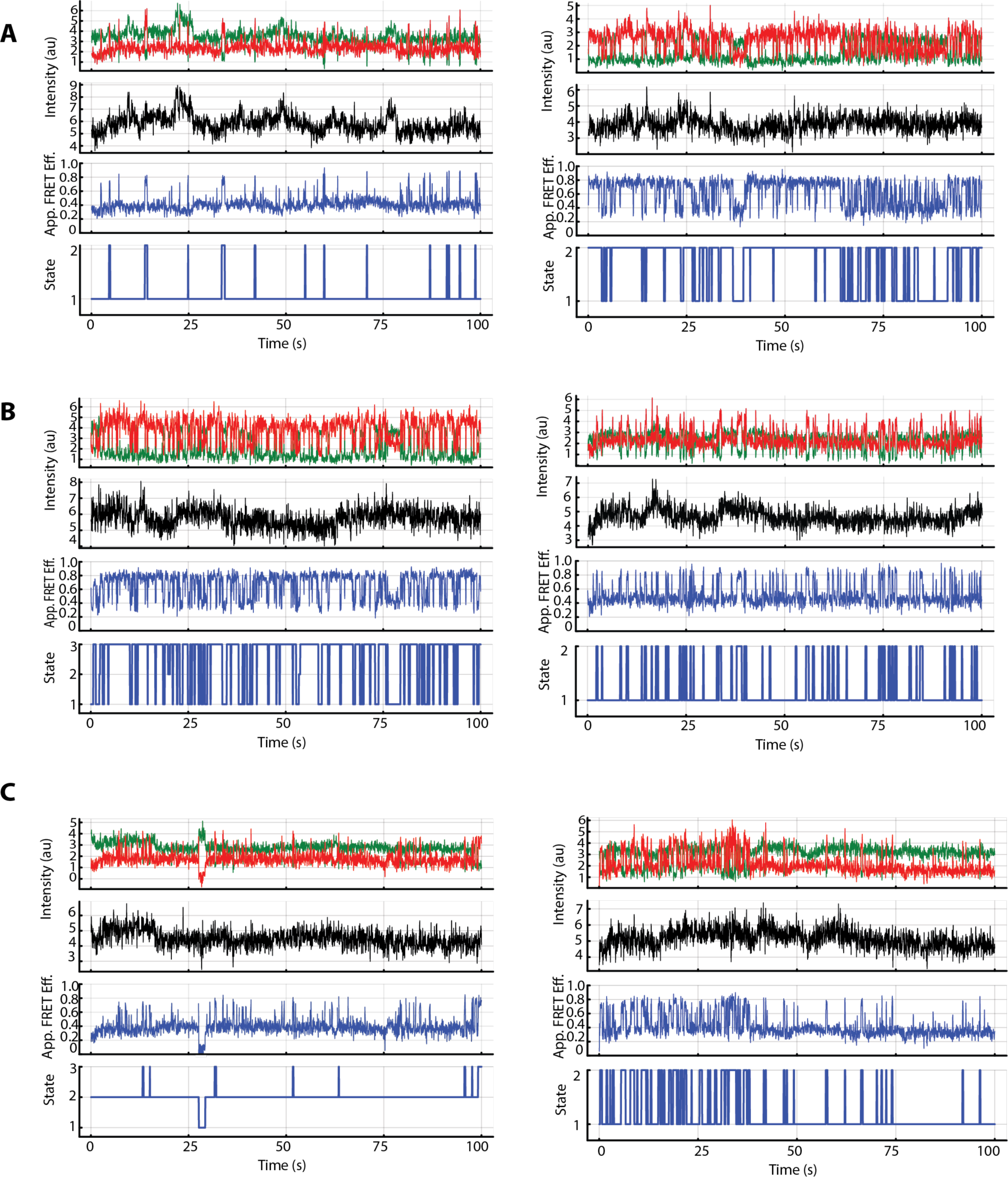
Representative traces from the conformational dynamics assay monitoring reconstituted wild-type **(A)**, YA6 **(B)**, and YA4-mutant proteasomes **(C)** in the absence of substrate. The first panels show fluorescence intensities for the FRET donor (green) and acceptor (red), the second panels depict the total fluorescence intensities (black), the third panels show the apparent FRET efficiencies, and the fourth panels illustrate the estimated most-probable state trajectories.

**Supp. Fig. 3-2.**
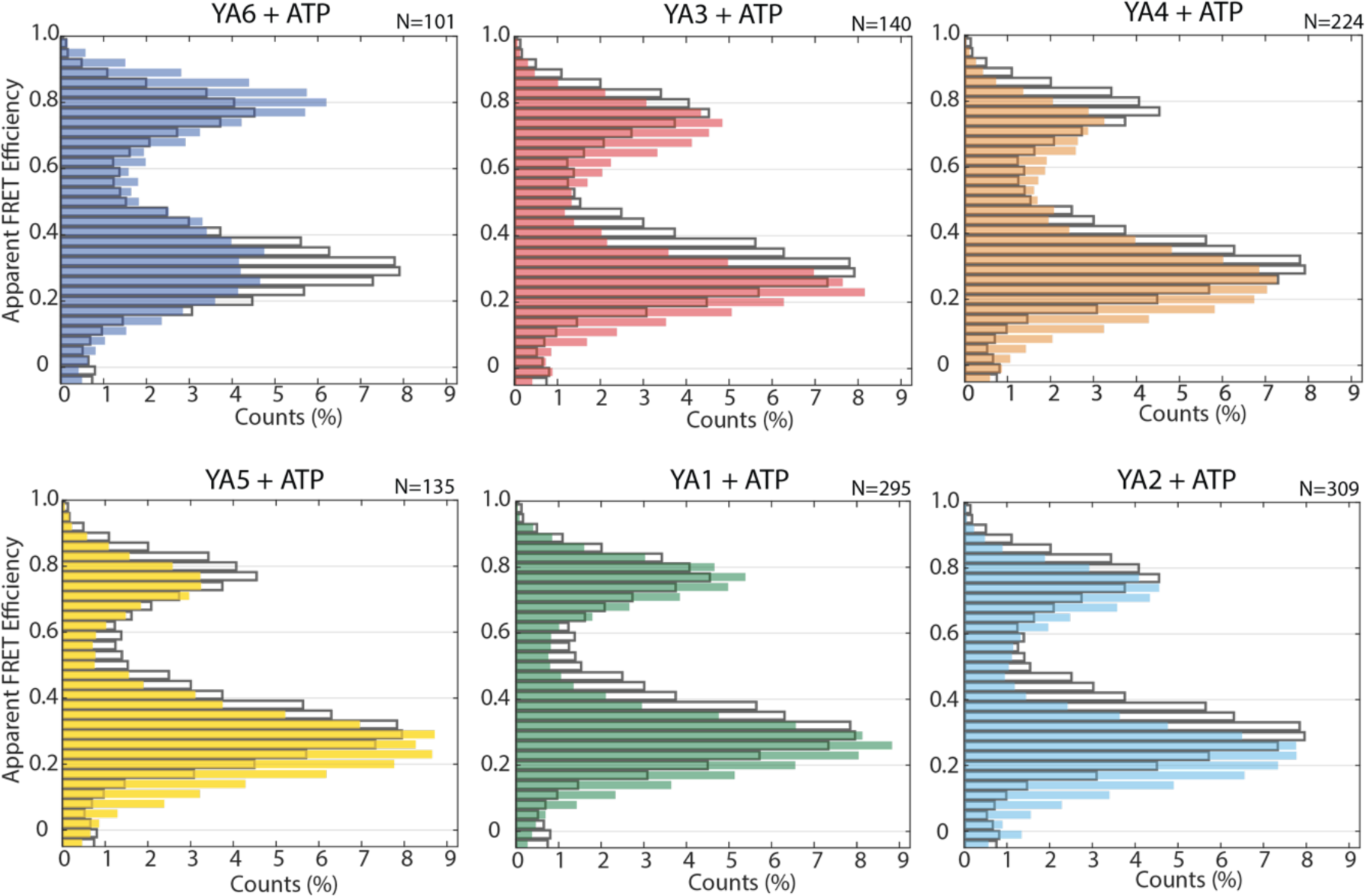
Apparent FRET efficiency distributions for YA6 (dark blue), YA3 (red),YA4 (orange), YA5 (yellow), YA1 (green), and YA2 (light blue) mutant proteasomes in ATP compared to the distribution of the wild-type proteasome (grey outlined bars).

**Supp. Fig. 3-3.**
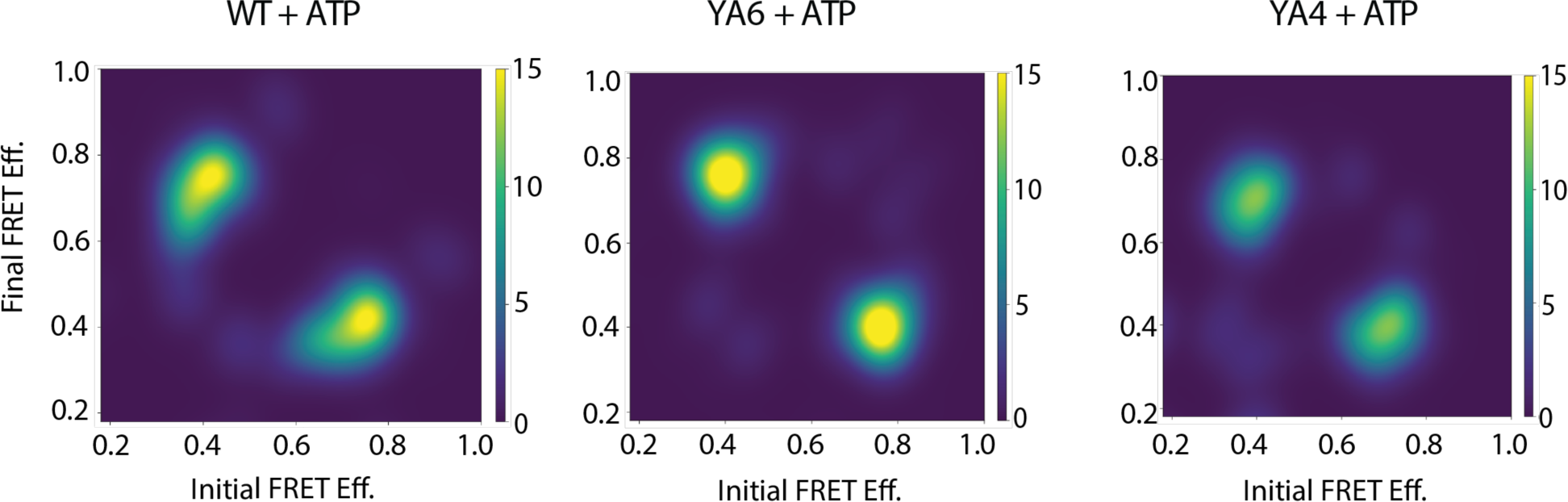
Transition density plots for wild-type, YA6, and YA4-muant 26S proteasomes in the absence of substrate. A scale color map was used (side bar), indicating the density of transitions within the states sampled.

**Supp. Fig. 4-1.**
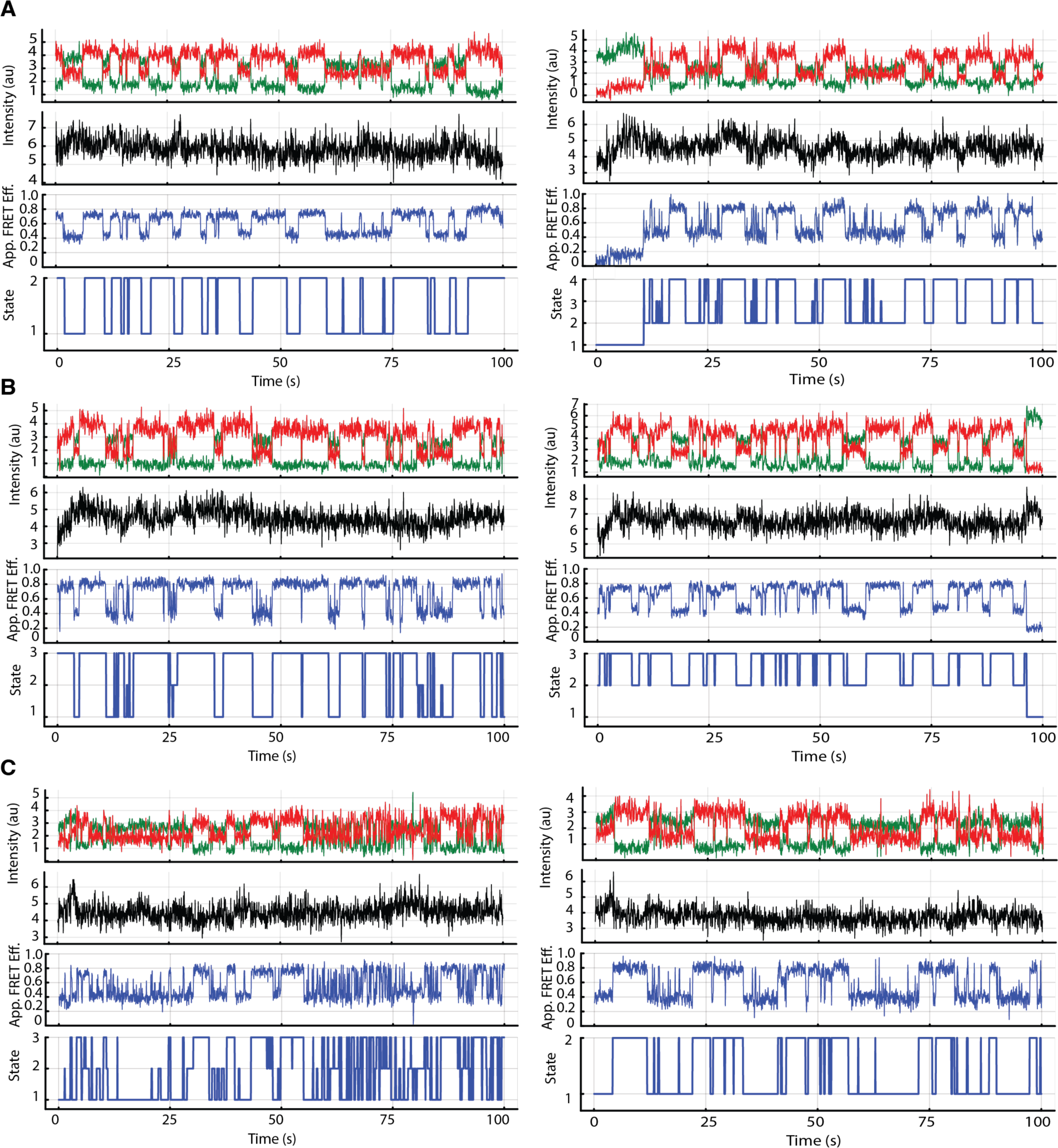
Representative traces from the conformational dynamics assay monitoring reconstituted wild-type **(A)**, YA6 **(B)**, and YA4-mutant proteasomes **(C)** in the presence of 1 μM titin I27^V15P^ substrate. The first panels show fluorescence intensities for the FRET donor (green) and acceptor (red), the second panels depict the total fluorescence intensities (black), the third panels show the apparent FRET efficiencies, and the fourth panels illustrate the estimated most-probable state trajectories.

**Supp. Fig. 4-2.**
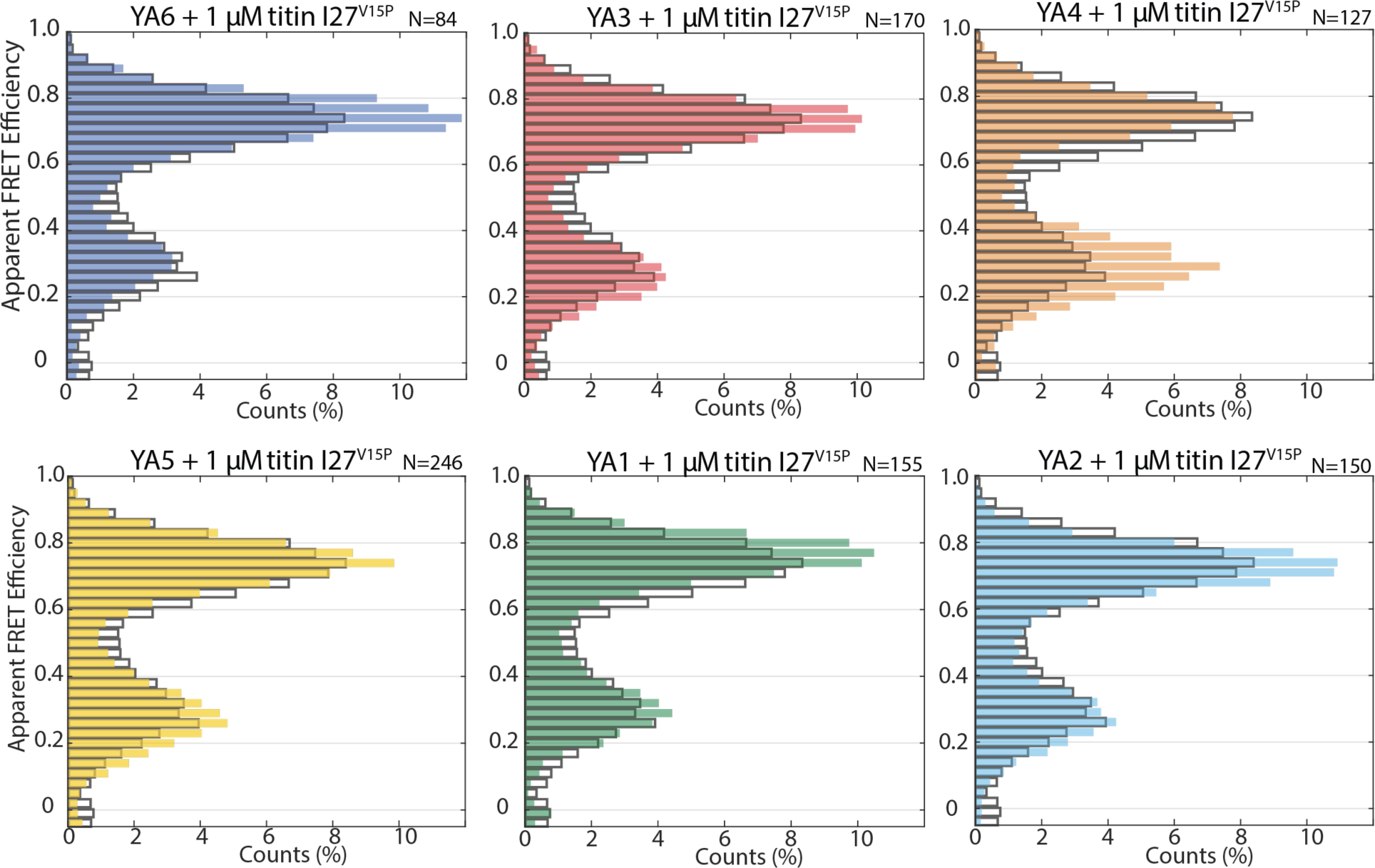
Apparent FRET efficiency distributions for YA6 (dark blue), YA3 (red),YA4 (orange), YA5 (yellow), YA1 (green), and YA2 (light blue) mutant proteasomes in the presence of 1 μM titin I27^V15P^ substrate compared to the distribution of the wild-type proteasome (grey outlined bars).

**Supp. Fig. 4-3.**
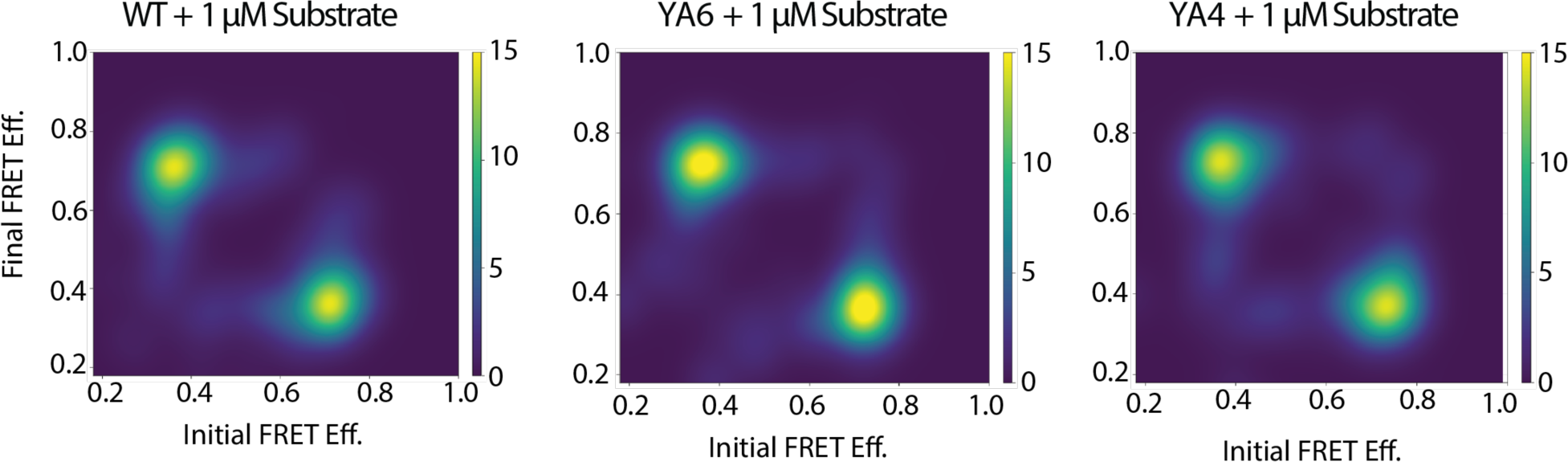
Transition density plots for wild-type, YA6, and YA4-muant 26S proteasomes in the presence of 1 μM titin I27^V15P^ substrate. A scale color map was used (side bar), indicating the density of transitions within the states sampled.

**Supp. Fig. 5-1.**
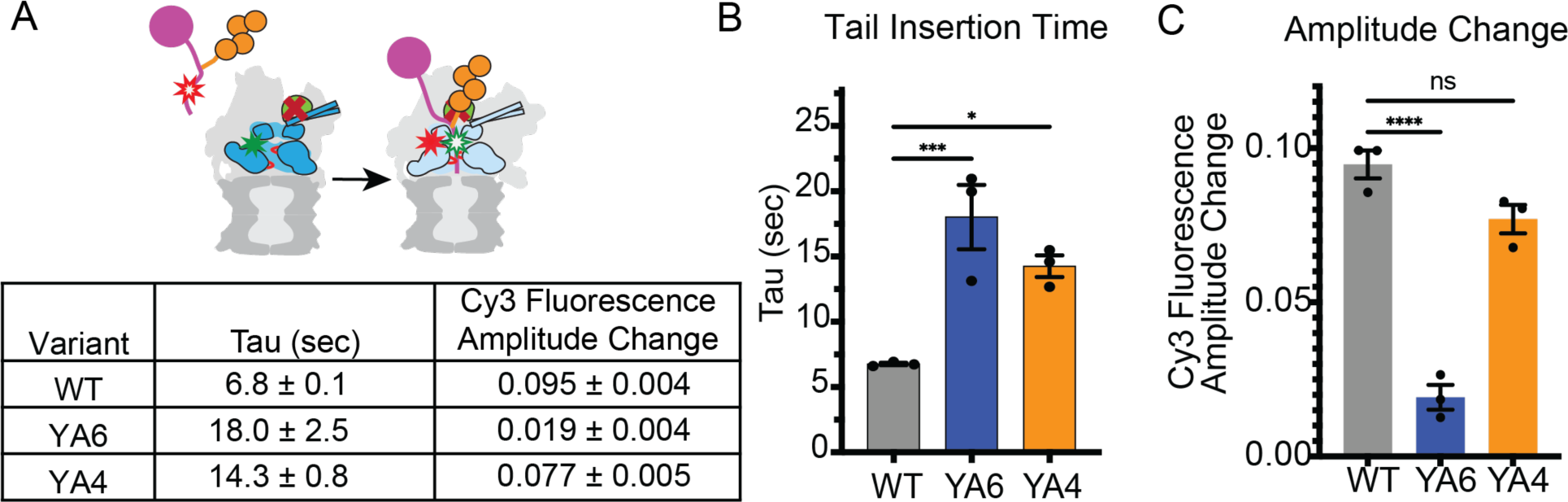
**A)** Top: Schematic for the ensemble measurements of substrate-tail insertion by monitoring the FRET efficiency between SulfoCy3-labeled 26S proteasomes with inhibited Rpn11 deubiquitinase (indicated by a red cross) and SulfoCy5-labeled substrate. The substrate and reconstituted proteasomes were rapidly mixed in a stopped flow instrument. Bottom: Time constants and Cy3-fluorescence amplitude changes (AU) during substrate tail insertion and engagement by wild-type and YA-mutant proteasomes. The error represents the SEM. **B)** Plotted time constants for tail insertion and **C)** Cy3-fluorescence amplitude changes obtained from the single exponential fitting of curves shown in Fig. 5B (N ≥ 3, technical replicates, error bars represent the SEM). Statistical significance was calculated using a one-way ANOVA test: ****p < 0.0001; **p < 0.01; *p < 0.05; ns, p > 0.05.

**Supp. Fig. 5-2.**
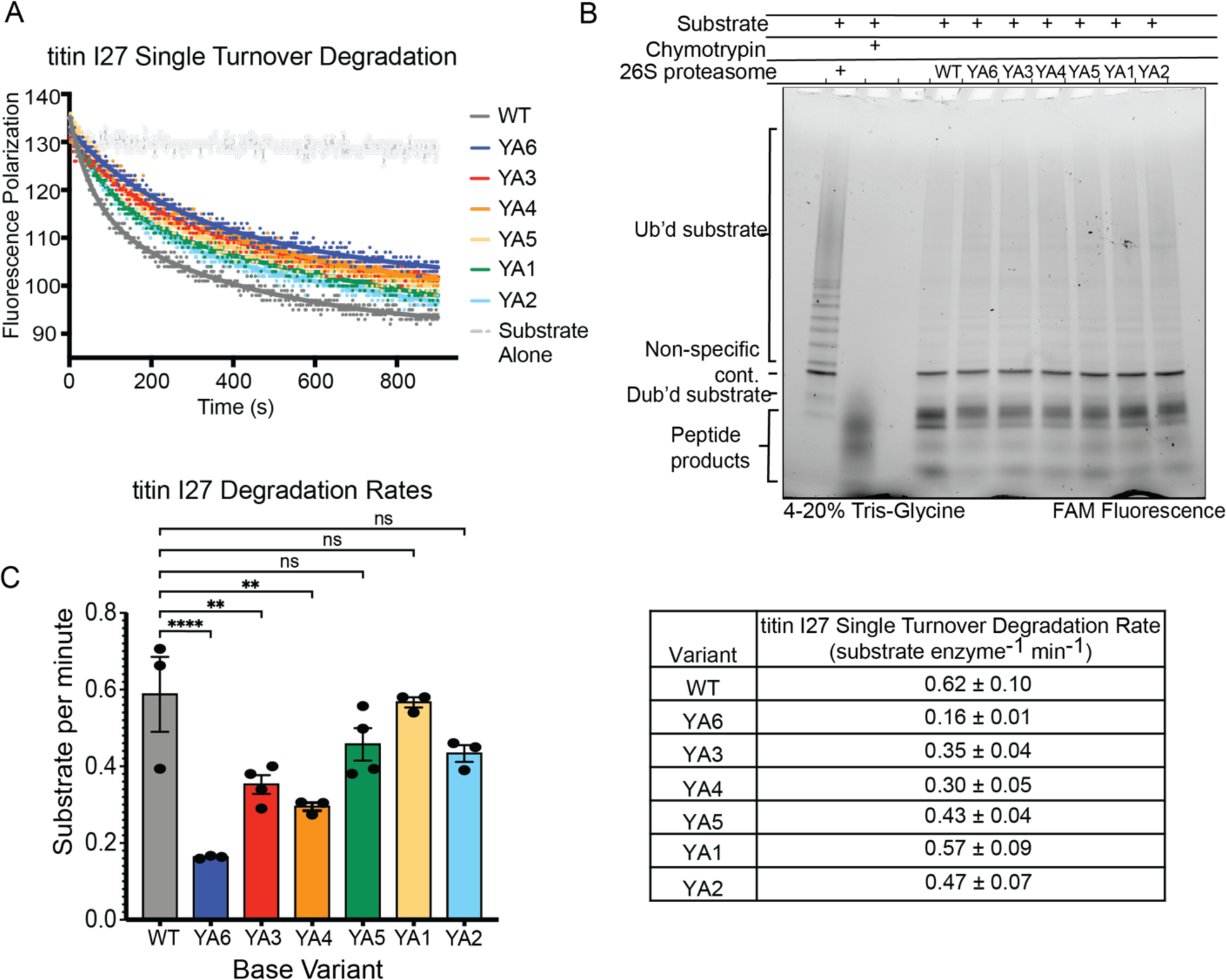
**A)** Representative traces for the single-turnover degradation of ubiquitinated FAM-titin I27 substrate by *in vitro*-reconstituted proteasomes. **B)** SDS-PAGE visualization of end- point samples from the single-turnover degradation of the FAM-titin I27 substrate by *in vitro*-reconstituted proteasomes. **C)** Single-turnover degradation rates obtained from fitting the curves shown in panel (A) to a double exponential decay using GraphPad Prism. (N = 3, technical replicates, error bars represent the SEM). Statistical significance was calculated using a one-way ANOVA test: ****p < 0.0001;**p < 0.01; ns, p > 0.05.

**Supp. Fig. 5-3.**
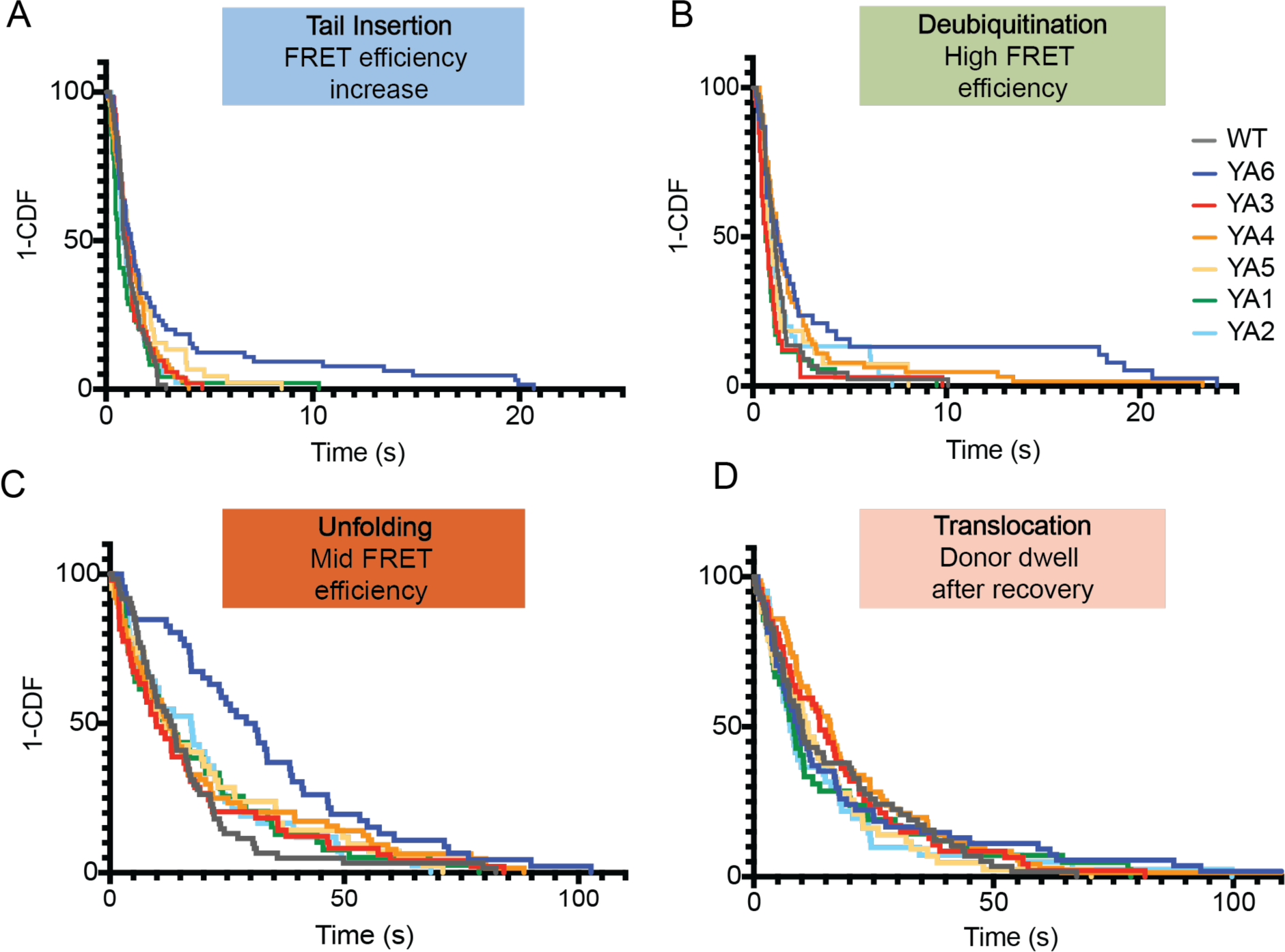
Single-molecule FRET-based measurements of titin I27 substrate processing by wild-type and YA-mutant proteasomes. 1-CDF (1-cumulative distribution function) or survival plot analyses for **A)** substrate tail insertion, **B)** deubiquitination, **C)** unfolding, and **D)** translocation after unfolding.

**Supp. Fig. 5-4.**
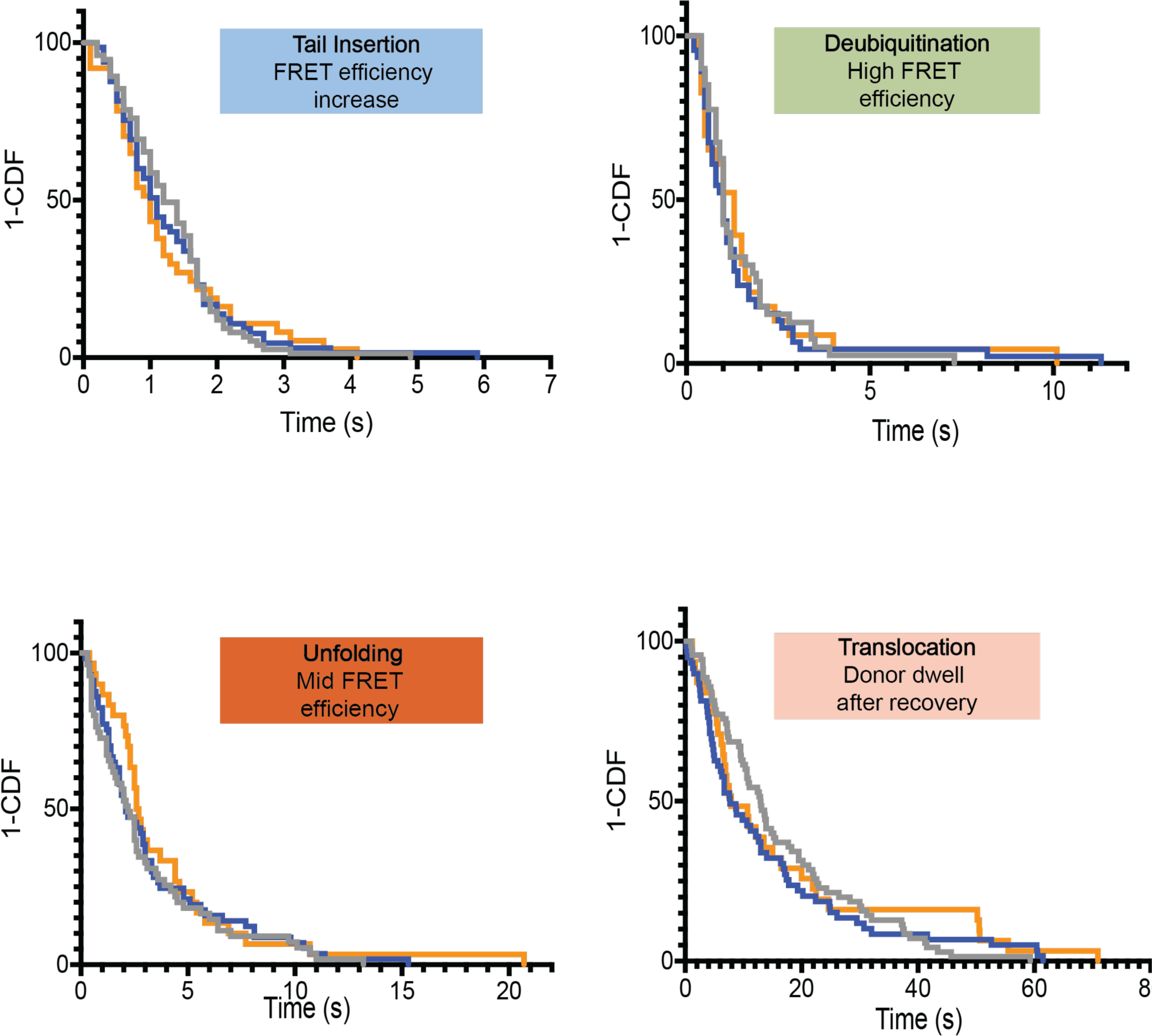
Single-molecule FRET-based measurements of titin I27^V15P^ substrate processing by wild-type and YA-mutant proteasomes. 1-CDF (1-cumulative distribution function) or survival plot analyses for **A)** substrate tail insertion, **B)** deubiquitination, **C)** unfolding, and **D)** translocation after unfolding.

**Supp. Fig. 5-5.**
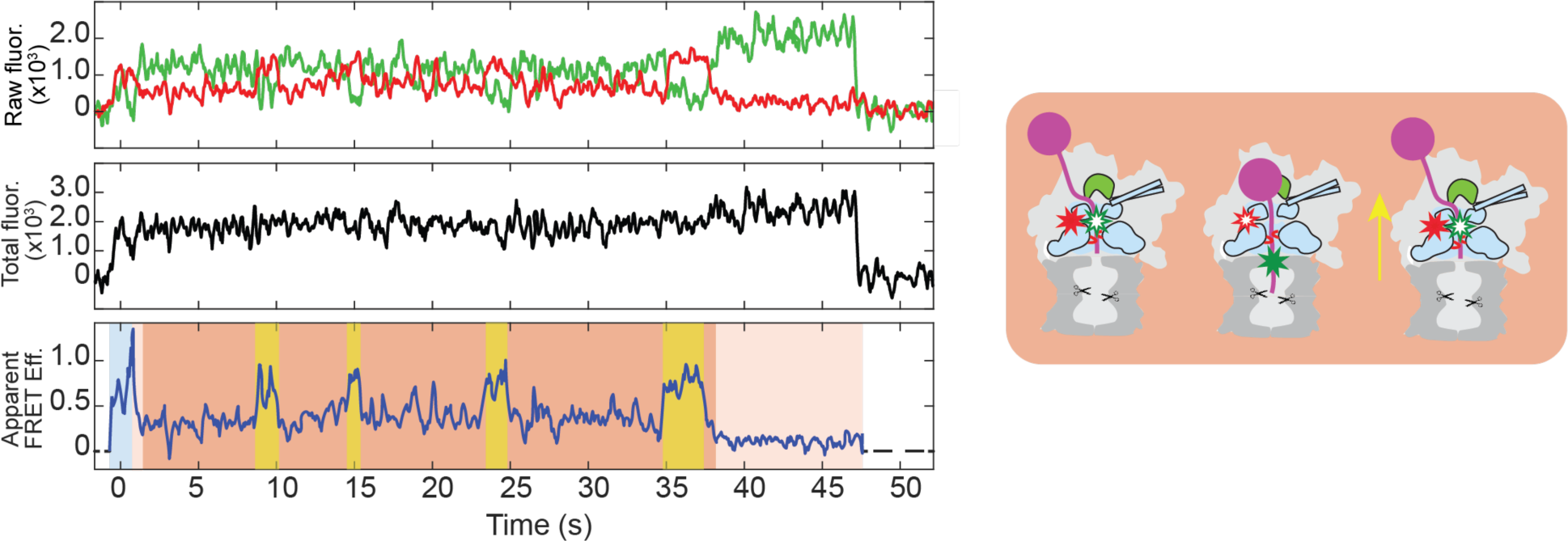
Backsliding events (yellow shading) of the titin I27 substrate during the unfolding phase (orange shading) by the YA3-mutant proteasomes, observed by single-molecule FRET.

**Supp. Fig. 6-1.**
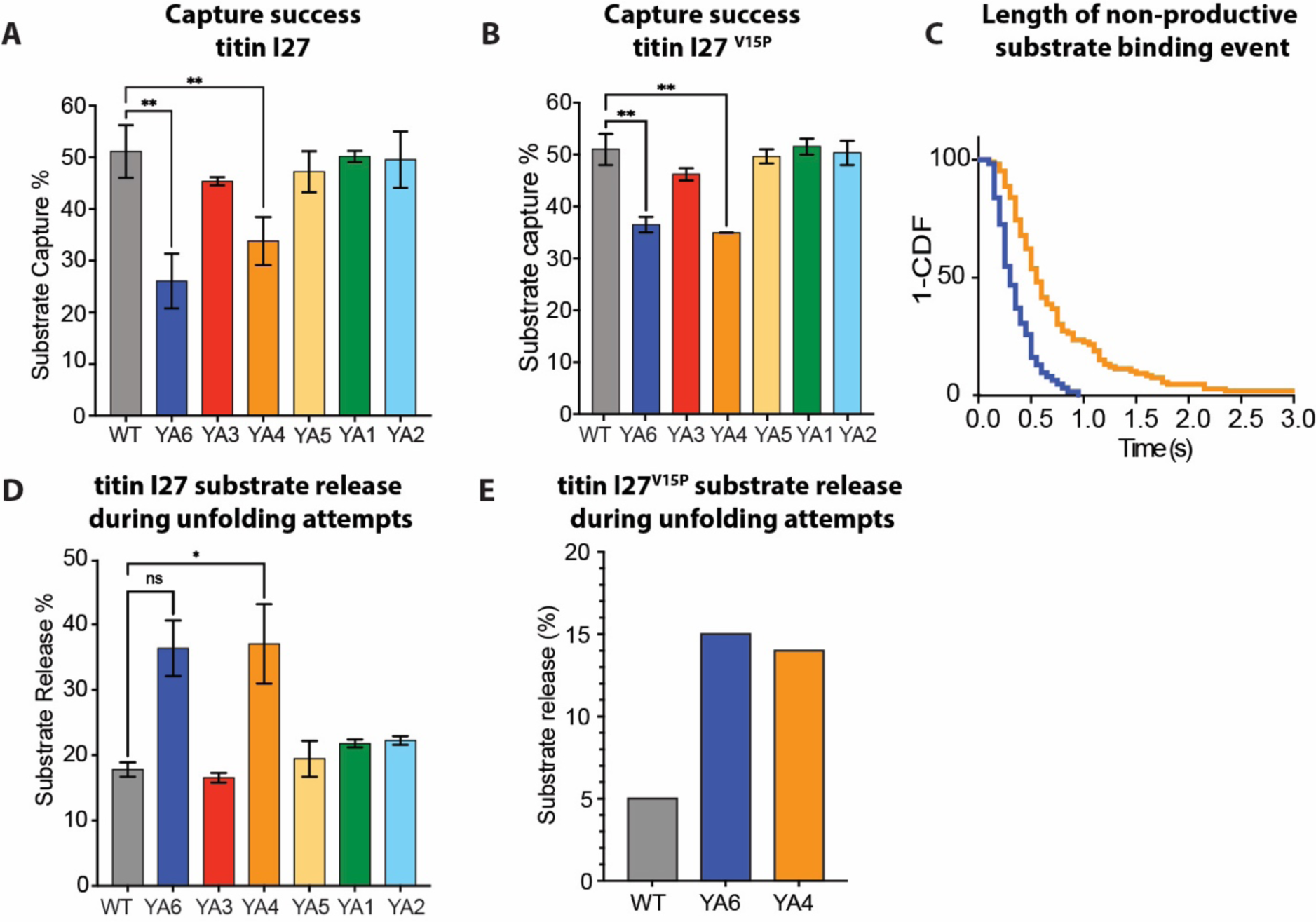
**A)** Success rates for the capture of wild-type titin I27 **(A)** and titin I27^V15P^ substrate **(B)** by wild-type and YA-mutant proteasomes. For these calculation, N = 2 technical replicates were obtained, contributing to n > 160 events for wild-type titin I27 and n > 150 for titin I27^V15P^. Error bars represent the SD, and statistical significance was calculated using a one-way ANOVA test: **p<0.01; *p<0.05; ns, p>0.05. **C)** Survival plots for brief events of substrate binding to YA4 and YA6-mutant proteasomes that do not lead to successful degradation. (N= 106 for YA4, N=62 for YA6) **D)** Percent of wild-type titin I27 substrate release during unfolding attempts by wild-type and YA-mutant proteasomes. N ≥ 2 technical replicates were performed, contributing the following number of events: WT = 103, YA6 = 110, YA3 = 90, YA4 = 142, YA5 = 75, YA1 = 83, YA2 = 81. **E)** Percent of titin I27^V15P^ substrate release during unfolding attempts by wild-type, YA6-, and YA4-mutant proteasomes. N ≥ 1 technical replicates were measured, contributing the following number of events: WT = 74, YA6 = 64, YA4 = 37.

**Supp. Fig. 6-2.**
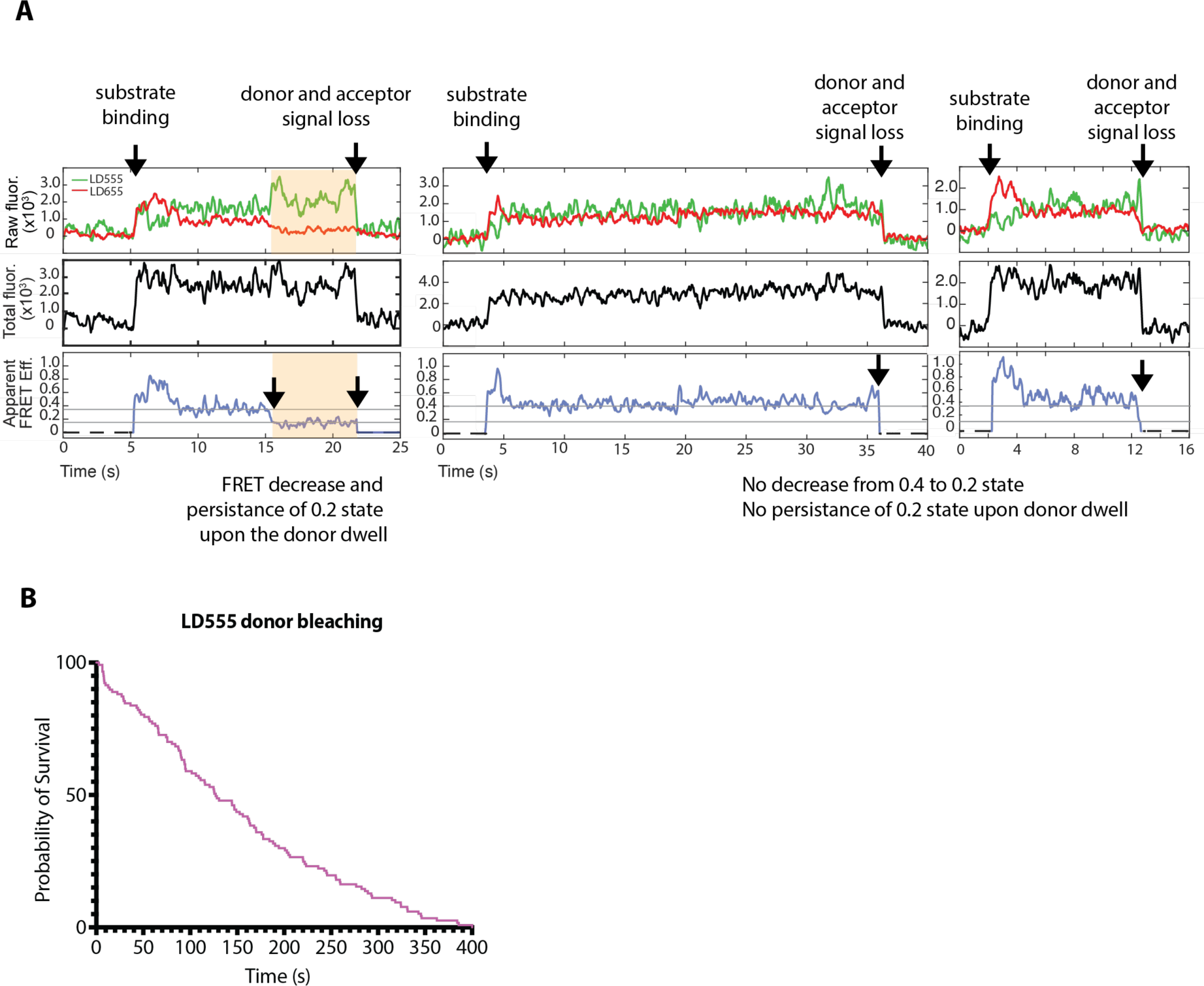
Substrate-release calculations. **A)** Substrate degradation events were scored as previously described. Traces on the left show a successful degradation event, with the FRET efficiency decreasing from ∼ 0.40 to ∼ 0.20 (signifying a progression from the pre-unfolding dwell to translocation of the unfolded protein) and then persisting at ∼ 0.2 during translocation. Substrate release (middle and right traces) was evaluated as the abrupt loss of donor and acceptor signal during the unfolding dwell, i.e. at the ∼ 0.40 FRET-efficiency state, with no decrease in FRET signal from ∼ 0.40 to ∼ 0.20 or a persistent 0.2 FRET efficiency. **B)** Measurement of the LD555 donor life time using LD555-labeled proteasomes in the absence of acceptor. A total of 152 particles were analyzed, out of which 26 % (n = 39) did not show donor photobleaching within the monitored time of 400 s. The remaining 78 % (n = 113) of photobleaching events were plotted to calculate a t_1/2_ ∼ 127 s.

## Supplemental Tables

**Supplementary Table 1:** Mean transition rates of proteasome conformational switching determined by HMM-MCMC. Traces resulting from the FRET-based conformational change assay for wild-type and pore-1 loop mutant proteasomes in the absence and presence of ubiquitinated titin I27^V15P^ were analyzed using MCMC-HMM techniques to determine the rates for the conformational transitions from the s1 to non-s1 states (k_s1_) and from non-s1 to the s1 state (k_non-s1_).

| 26S proteasome sample | Mean $k_{s1}$ ($s^{-1}$) | Mean $k_{non-s1}$ ($s^{-1}$) |
| --- | --- | --- |
| WT + 2 mM ATP | 1.30 | 1.93 |
| YA6 + 2 mM ATP | 1.57 | 2.25 |
| YA4 + 2 mM ATP | 0.99 | 2.03 |
| WT + 2 mM ATP + 1 μM titin I27<sup>V15P</sup> | 1.72 | 0.54 |
| YA6 + 2 mM ATP + 1 μM titin I27<sup>V15P</sup> | 1.43 | 0.63 |
| YA4 + 2 mM ATP + 1 μM titin I27<sup>V15P</sup> | 1.18 | 1.05 |

**Supplementary Table 2.**
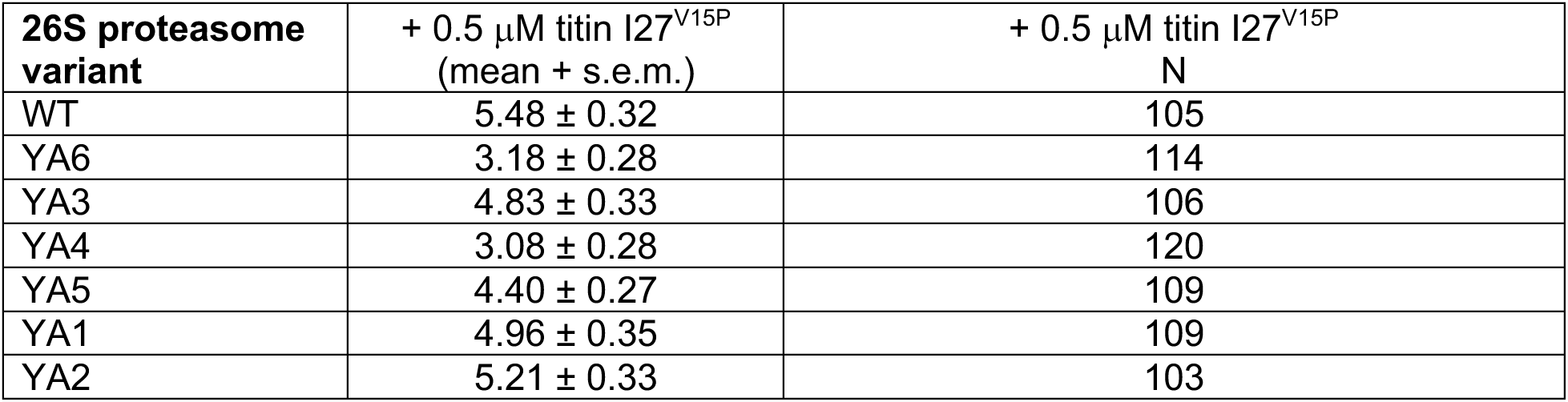
Substrate processing dwells. Listed are the mean values for the duration of high-FRET processing dwells during the degradation of 0.5 μM ubiquitinated titin I27^V15P^ as measured by the FRET-based conformational change assay. The N values indicate the number of analyzed persistent high-FRET events.

| 26S proteasome variant | + 0.5 μM titin I27<sup>V15P</sup><br>(mean + s.e.m.) | + 0.5 μM titin I27<sup>V15P</sup><br>N |
| --- | --- | --- |
| WT | 5.48 ± 0.32 | 105 |
| YA6 | 3.18 ± 0.28 | 114 |
| YA3 | 4.83 ± 0.33 | 106 |
| YA4 | 3.08 ± 0.28 | 120 |
| YA5 | 4.40 ± 0.27 | 109 |
| YA1 | 4.96 ± 0.35 | 109 |
| YA2 | 5.21 ± 0.33 | 103 |

**Supplementary Table 3.** Titin I27 tail insertion and engagement kinetics. Listed are the time constants for tail insertion and engagement of ubiquitinated titin I27 substrate, as determined by fitting the survival plots for the tail-insertion phase in the FRET-based processing assay to a single exponential. Given are also the values for a 95% confidence interval (CI), and the N values indicate the number of analyzed degradation events.

| 26S proteasome variant | Tau (s) | 95% CI | N |
| --- | --- | --- | --- |
| WT | 1.48 | 1.20 - 1.90 | 65 |
| YA6 | 1.42 | 1.29 - 1.55 | 66 |
| YA3 | 1.22 | 1.00 - 1.52 | 52 |
| YA4 | 1.63 | 1.47 - 1.83 | 72 |
| YA5 | 1.47 | 1.34 - 1.61 | 45 |
| YA1 | 0.79 | 0.69 - 0.91 | 50 |
| YA2 | 1.06 | 0.92 - 1.23 | 47 |

**Supplementary Table 4.** Titin I27 deubiquitination kinetics. Listed are the time constants for deubiquitination of ubiquitinated titin I27 substrate, as determined by fitting the survival plots for the high-FRET deubiquitination phase in the FRET-based processing assay to a single exponential. Given are also the values for a 95% confidence interval (CI), and the N values indicate the number of analyzed degradation events.

| **26S proteasome variant** | **Tau (s)** | **95% CI** | **N** |
| --- | --- | --- | --- |
| WT | 1.29 | 1.09 - 1.54 | 44 |
| YA6 | 1.49 | 1.37 - 1.63 | 39 |
| YA3 | 0.85 | 0.73 - 0.98 | 33 |
| YA4 | 1.51 | 1.39 - 1.63 | 64 |
| YA5 | 1.03 | 0.84 - 1.29 | 27 |
| YA1 | 0.81 | 0.65 - 1.03 | 36 |
| YA2 | 1.10 | 0.86 - 1.19 | 30 |

**Supplementary Table 5.** Titin I27 unfolding times. Listed are the time constants for overall successful unfolding attempts (pre-unfolding dwells) during the degradation of ubiquitinated titin I27 substrate, as determined by fitting the survival plots for the unfolding phase in the FRET-based processing assay to a single exponential. Given are also the values for a 95% confidence interval (CI), and the N values indicate the number of analyzed degradation events.

| **26S proteasome variant** | **Tau (s)** | **95% CI** | **N** |
| --- | --- | --- | --- |
| WT | 16.31 | 14.96 - 17.82 | 61 |
| YA6 | 50.82 | 43.50 - 60.58 | 46 |
| YA3 | 13.46 | 12.51 - 14.49 | 52 |
| YA4 | 15.43 | 14.53 - 16.40 | 64 |
| YA5 | 18.10 | 16.56 - 19.85 | 45 |
| YA1 | 17.29 | 15.21 - 19.73 | 47 |
| YA2 | 21.06 | 19.24 - 23.14 | 47 |

**Supplementary Table 6.**
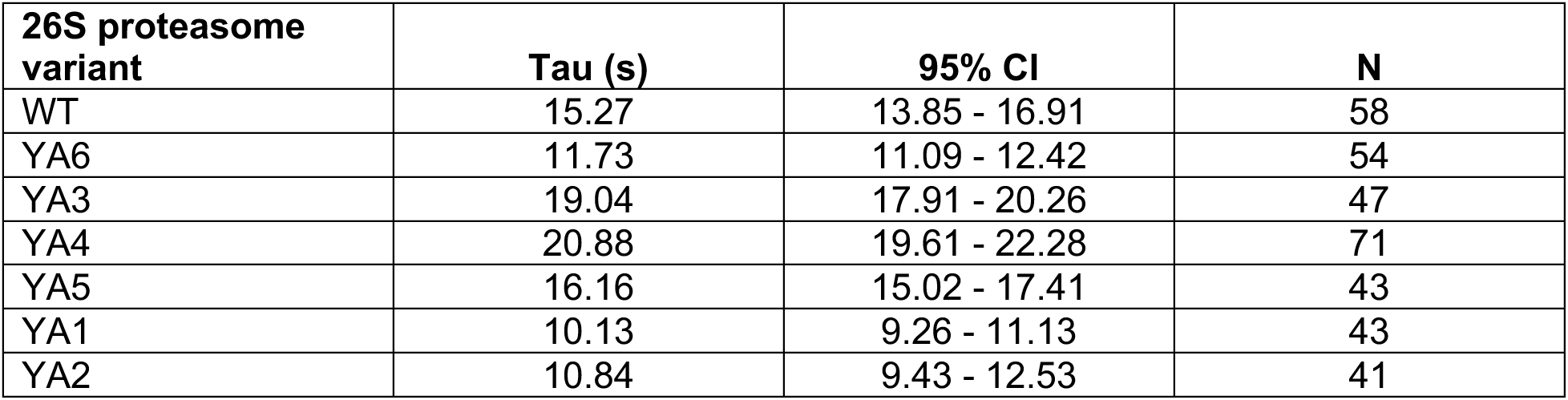
Titin I27 translocation kinetics. Listed are the time constants for the translocation of titin I27 substrate after deubiquitination and unfolding, as determined by fitting the survival plots for the donor dwell after donor-fluorescence recovery in the FRET-based processing assay to a single exponential. Given are also the values for a 95% confidence interval (CI), and the N values indicate the number of analyzed degradation events.

| **26S proteasome variant** | **Tau (s)** | **95% CI** | **N** |
| --- | --- | --- | --- |
| WT | 15.27 | 13.85 - 16.91 | 58 |
| YA6 | 11.73 | 11.09 - 12.42 | 54 |
| YA3 | 19.04 | 17.91 - 20.26 | 47 |
| YA4 | 20.88 | 19.61 - 22.28 | 71 |
| YA5 | 16.16 | 15.02 - 17.41 | 43 |
| YA1 | 10.13 | 9.26 - 11.13 | 43 |
| YA2 | 10.84 | 9.43 - 12.53 | 41 |

**Supplementary Table 7.** Titin I27^V15P^ substrate processing kinetics. Listed are the time constant values for tail insertion, deubiquitination, pre-unfolding dwell, and translocation of the ubiquitinated titin I27^V15P^ substrate, as determined by single exponential decay fitting of the FRET-based tail insertion phase length in the substrate processing assay. In parenthesis are the values for the 95% confidence interval. N values indicate the number of analyzed events.

| 26S proteasome variant | Tail Insertion |  | Deubiquitination |  | Pre-unfolding |  | Translocation |  |
| --- | --- | --- | --- | --- | --- | --- | --- | --- |
|  | Tau (s) | N | Tau (s) | N | Tau (s) | N | Tau (s) | N |
| WT | 1.88<br>(1.45 - 2.58) | 69 | 1.49<br>(1.13 - 2.03) | 32 | 2.76<br>(2.57 - 2.97) | 60 | 18.46<br>(17.27 - 19.80) | 70 |
| YA6 | 1.50<br>(1.25 - 1.81) | 72 | 1.20<br>(1.00 - 1.45) | 35 | 2.77<br>(2.58 - 2.98) | 61 | 11.71<br>(11.05 - 12.42) | 71 |
| YA4 | 1.37<br>(1.09 - 1.79) | 67 | 1.41<br>(1.09 - 1.85) | 34 | 3.73<br>(3.06 - 4.61) | 62 | 11.91<br>(10.48 - 13.61) | 51 |

**Supplementary Table 8.**
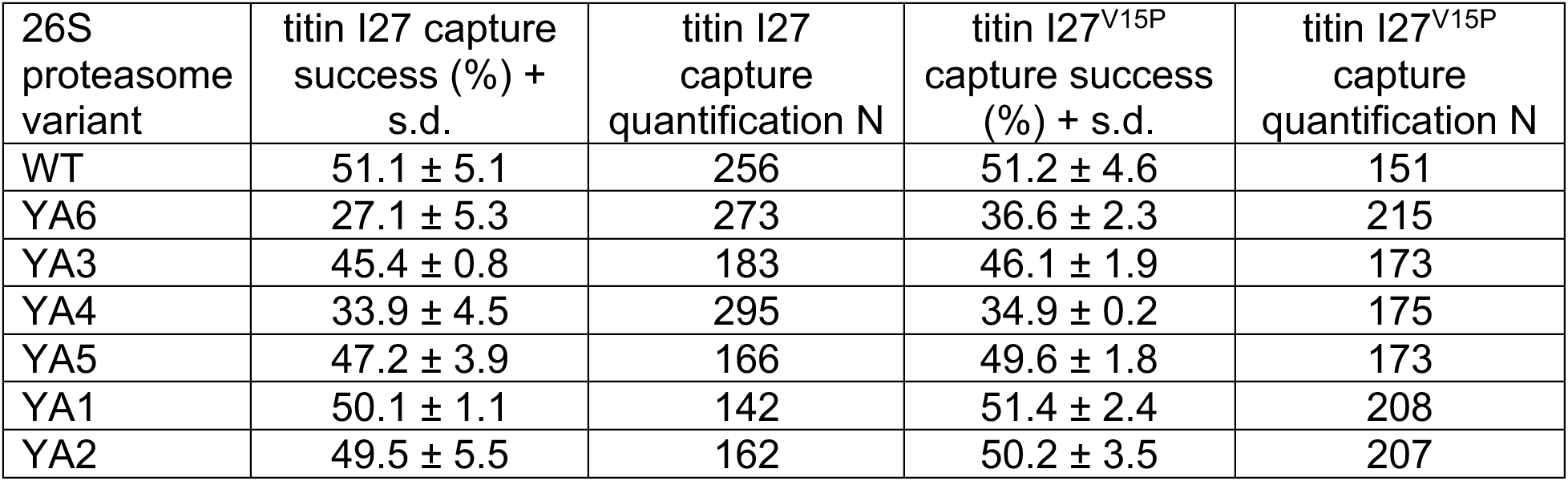
Substrate capture success by 26S proteasome variants. Listed are the average values and standard deviation for substrate capture success percentage of the ubiquitinated titin I27 and titin I27^V15P^ substrates. N values indicate the number of analyzed events.

**Supplementary Table 9.**
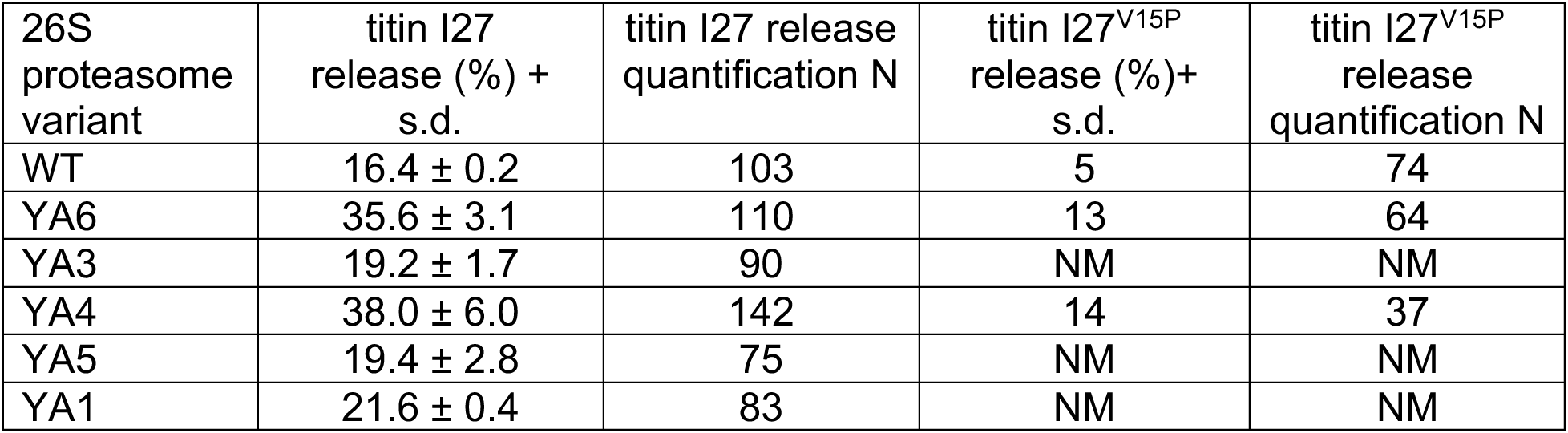
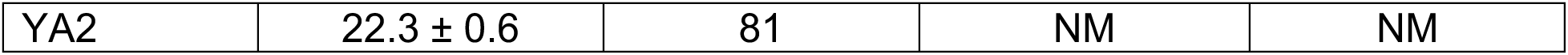
Substrate release upon pre-unfolding dwell 26S proteasome variants. Listed are the average values and standard deviation for substrate release percentage of the ubiquitinated titin I27 and titin I27^V15P^ substrates. N values indicate the number of analyzed events. NM – not measured

**Supplementary Table 10.** Percentage of particles from different cryo-EM datasets in which the Rpt4 subunit adopts the bottom or seam position within the spiral staircase arrangement of the AAA+ motor. The percentage calculated is relative to the total number of particles observed in processing-competent states. For these studies, yeast or human 26S proteasome were incubated with a substrate and stalled through different methods, either the deubiquitinase inhibitor 1,10-phenanthroline or a mixture of ATP and the non-hydrolysable ATP analog ATPγS. Given in parentheses in the third column are the respective processing states based on the specific nomenclature of the authors.

| Study | Sample | Reported # of particles with Rpt4 in the bottom or seam position (state according to author nomenclature) |
| --- | --- | --- |
| De la Peña *et al.*, 2018<sup>31</sup> | *S. cerevisiae* 26S<br>+ Ub'd titin I27<sup>V15P</sup><br>+ DUB inhibitor<br>+ ATP | 30% (4D)<br>29% (5T)<br>19.2% (5D) |
| Dong *et al.*, 2018<sup>42</sup> | Human 26S<br>+ Ub'd-Sic1PY<br>+ ATP/ATP<sub>γ</sub>S | 60% (E<sub>D2</sub>)<br>40% (E<sub>D1</sub>) |
| Zhang *et al.*, 2022<sup>37</sup> | Human 26S<br>+ Ub-Sic1PY<br>+ Usp14<br>+ ATP/ATP<sub>γ</sub>S | 64% (E<sub>D2</sub> E<sub>D2.1</sub>)<br>12% (E<sub>D1</sub>) |

**Supplementary Table 11.** Plasmids utilized in this study.

| Plasmid ID | Phenotype | Source |
| --- | --- | --- |
| pAM371 | pCOLA FLAG-Rpt1-Y283A, Rpt2, His<sub>6</sub>-Rpt3, Rpt5, Rpt6, Rpt4 | This study (derived from Beckwith *et al.*, 2013) |
| pAM372 | pCOLA FLAG-Rpt1, Rpt2-Y256A, His<sub>6</sub>-Rpt3, Rpt5, Rpt6, Rpt4 | This study (derived from Beckwith *et al.*, 2013) |
| pAM373 | pCOLA FLAG-Rpt1, Rpt2, His<sub>6</sub>-Rpt3-Y246A, Rpt5, Rpt6, Rpt4 | This study (derived from Beckwith *et al.*, 2013) |
| pAM374 | pCOLA FLAG-Rpt1, Rpt2, His<sub>6</sub>-Rpt3, Rpt5, Rpt6, Rpt4-Y256A | This study (derived from Beckwith *et al.*, 2013) |
| pAM375 | pCOLA FLAG-Rpt1, Rpt2, His<sub>6</sub>-Rpt3, Rpt5-Y255A, Rpt6, Rpt4 | This study (derived from Beckwith *et al.*, 2013) |
| pAM376 | pCOLA FLAG-Rpt1, Rpt2, His<sub>6</sub>-Rpt3, Rpt5, Rpt6-Y222A, Rpt4 | This study (derived from Beckwith *et al.*, 2013) |
| pAM82 | pCOLA FLAG-Rpt1, Rpt2, His<sub>6</sub>-Rpt3, Rpt5, Rpt6, Rpt4 | Bard *et al.*, 2019 |
| pAM377 | pCOLA FLAG-Rpt1-I191TAG-Y283A, Rpt2, His<sub>6</sub>-Rpt3, Rpt5, Rpt6, Rpt4 | This study |
| pAM378 | pCOLA FLAG-Rpt1-I191TAG, Rpt2-Y256A, His<sub>6</sub>-Rpt3, Rpt5, Rpt6, Rpt4 | This study |
| pAM379 | pCOLA FLAG-Rpt1-I191TAG, Rpt2, His<sub>6</sub>-Rpt3-Y246A, Rpt5, Rpt6, Rpt4 | This study |
| pAM380 | pCOLA FLAG-Rpt1-I191TAG, Rpt2, His<sub>6</sub>-Rpt3, Rpt5, Rpt6, Rpt4-Y256A | This study |
| pAM381 | pCOLA FLAG-Rpt1-I191TAG, Rpt2, His<sub>6</sub>-Rpt3, Rpt5-Y255A, Rpt6, Rpt4 | This study |
| pAM382 | pCOLA FLAG-Rpt1-I191TAG, Rpt2, His<sub>6</sub>-Rpt3, Rpt5, Rpt6-Y222A, Rpt4 | This study |
| pAM88 | pCOLA FLAG-Rpt1-I191TAG, Rpt2, His<sub>6</sub>-Rpt3, Rpt5, Rpt6, Rpt4 | Bard et al., 2019 |
| pAM383 | pCOLA FLAG-Rpt1-Y283A, Rpt2, His<sub>6</sub>-Rpt3, Rpt5-Q49TAG, Rpt6, Rpt4 | This study |
| pAM384 | pCOLA FLAG-Rpt1, Rpt2-Y256A, His<sub>6</sub>-Rpt3, Rpt5-Q49TAG, Rpt6, Rpt4 | This study |
| pAM385 | pCOLA FLAG-Rpt1, Rpt2, His<sub>6</sub>-Rpt3-Y246A, Rpt5-Q49TAG, Rpt6, Rpt4 | This study |
| pAM386 | pCOLA FLAG-Rpt1, Rpt2, His<sub>6</sub>-Rpt3, Rpt5-Q49TAG, Rpt6, Rpt4-Y256A | This study |
| pAM387 | pCOLA FLAG-Rpt1, Rpt2, His<sub>6</sub>-Rpt3, Rpt5-Q49TAG-Y255A, Rpt6, Rpt4 | This study |
| pAM388 | pCOLA FLAG-Rpt1, Rpt2, His<sub>6</sub>-Rpt3, Rpt5-Q49TAG, Rpt6-Y222A, Rpt4 | This study |
| pAM89 | pCOLA FLAG-Rpt1, Rpt2, His<sub>6</sub>-Rpt3, Rpt5-Q49TAG, Rpt6, Rpt4 | Bard et al., 2019 |
| pAM81 | pETDuet1 Rpn1, Rpn2, Rpn13 | Beckwith et al., 2013 |
| pAM83 | pACYC-Duet RIL Nas6, Hsm3, Rpn14, Nas2 | Beckwith et al., 2013 |
| pAM87 | pUltra_pAzFRS.2.t1_UAG-tRNA | Bard et al., 2019 |
| pAM85 | pETDuet1 Rpn5, MBP-HRV3C-Rpn6, Rpn8, Rpn11, Rpn9 | Bard et al., 2019 |
| pAM314 | pETDuet1 Rpn5, MBP-HRV3C-Rpn6, Rpn8, Rpn11, Rpn9-F2TAG | Jonsson et al., 2022 |
| pAM80 | pACYC Sem1, Hsp90 | Lander et al, 2012 |
| pAM86 | pCOLA His<sub>6</sub>-HRV3C-Rpn12, Rpn7, Rpn3 | Bard et al., 2019 |
| pAM239 | pACYC Rpn10 | Beckwith et al., 2013 |
| pAM91 | Titin I27 V15P (lysineless)-PPPY-ssrA-1K-35 amino acid tail | Bard et al., 2019 |
| pAM93 | WT Titin I27 (lysineless)-PPPY-ssrA-1K-35 amino acid tail | Bard et al., 2019 |

**Supplementary Table 12.** Yeast strains utilized in this study.

| Yeast Strain | Phenotype / Expression | Source |
| --- | --- | --- |
| yAM54 | 3xFLAG-Pre1 | Beckwith et al., 2013 |
| yAM80 | Pre1-BirA-HRV-3xFLAG | Jonsson et al., 2022 |

## Materials and Methods

### Purification and labeling of base subcomplex

Recombinant base subcomplexes were expressed, purified, and labeled as previously described. For recombinant expression of the base subcomplex, *Escherichia coli* BL21-star DE3 cells were transformed with pAM81, pAM83, and the corresponding plasmid for Rpt1-6 subunits and pore-1 loop mutant expression (Supp. Table 11). For unnatural amino acid incorporation, an additional plasmids pAM87 was also co-transformed for expression of the AzF tRNA synthase/tRNA. Cells were cultured overnight in DYT media supplemented with 1x antibiotics (300 μg/mL ampicillin, 25 μg/mL chloramphenicol, 50 μg/mL kanamycin, and additional 100 μg/mL spectinomycin for unnatural amino acid incorporation). The next day, cells were grown at 37°C in 3 L of buffered terrific broth (17 mM KH_2_PO_4_ and 72 mM K_2_HPO_4_) media with 0.5x antibiotics to an optical density at 600 nm (OD_600nm_) of 0.8, then induced with 1 mM isopropyl-B-D-thiogalactopyranoside (IPTG) for 5 hours at 30°C and then at 16°C overnight. For unnatural amino acid incorporation, 2 mM AzF (Amatek Chemical) was added per liter before induction with IPTG. Cells were harvested by spinning at 3,500 rpm for 20 minutes at room temperature and then resuspended in lysis buffer (60 mM HEPES pH 7.60, 100 mM NaCl, 100 mM KCl, 10 mM MgCl_2_, 10% glycerol, and 20 mM imidazole) supplemented with protease inhibitors (aprotonin, leupeptin, pepstatin, and 4-(2-aminoethyl)benzenesulfonyl fluoride hydrochloride (AEBSF)), benzonase, and 2 mg/mL lysozyme.

The base subcomplex was purified in two affinity steps and one size exclusion step, with 1 mM ATP supplemented in all buffers. First, cells were lysed by sonication and centrifuged for 30 min at 15,000 rpm at 4°C for lysate clarification. The eluant was loaded to a 5 mL HisTrap FF crude (Cytiva) column using a peristaltic pump, washed with NiA buffer (60 mM HEPES pH 7.60, 100 mM NaCl, 100 mM KCl, 10 mM MgCl_2_, 10% glycerol, and 20 mM imidazole), and eluted with NiB buffer (60 mM HEPES pH 7.60, 100 mM NaCl, 100 mM KCl, 10 mM MgCl_2_, 10% glycerol, and 200 mM imidazole). The eluant was then flowed on a M2 Anti-FLAG affinity resin (Sigma Aldrich), washed with NiA buffer, and eluted with NiA buffer supplemented with 0.5 mg/mL 3xFLAG peptide. After concentrating with a 30 kDa molecular weight cut off (MWCO) concentrator (Millipore), the subcomplex was further purified by size-exclusion chromatography (SEC) using a Superose 6 increase 10/300 column (Cytiva) in GF buffer (30 mM HEPES pH 7.60, 50 mM NaCl, 50 mM KCl, 10 mM MgCl_2_, 5% glycerol) supplemented with 500 μM tris(2-carboxyethyl)phosphine (TCEP) and 1 mM ATP. For labeling via unnatural amino acid, 150 μM 5,50-dithiobis-2-nitrobenzoic acid (DTNB, Sigma Aldrich) was added to the purified base after concentrating and incubated for 10 min on ice before addition of 300 μM dybenzocyclooctane (DBCO)-Cy3 (Click Chemistry Tools) or LD655 (Lumidyne Technologies). The reaction progressed overnight at 4°C supplemented with ATP Regeneration Mix (creatine phosphate (VWR) and creatine kinase (Sigma Aldrich)) and was quenched the next day by incubating with 300 μM free AzF for 5 minutes and 5 mM DTT for 30 minutes at 4°C before SEC. The concentration of base subcomplex was determined by Bradford with a BSA (Sigma Aldrich) standard curve or and the labeling efficiency by quantification of the dye absorbance using an Agilent UV-Vis Spectrophotometer.

### Purification and labeling of lid subcomplex

Recombinant lid subcomplexes were expressed, purified, and labeled as previously described ^17,38^. For recombinant expression of the lid subcomplex, *Escherichia coli* BL21-star DE3 cells were transformed with pAM80, pAM86, and pAM85 (Supp. Table 11). For unnatural amino acid incorporation, pAM314 was used in place of pAM85 and an additional plasmid pAM87 was co-transformed for expression of the AzF tRNA synthase/tRNA. Cells were cultured overnight in DYT media supplemented with 1x antibiotics (300 μg/mL ampicillin, 25 μg/mL chloramphenicol, 50 μg/mL kanamycin, and additional 100 μg/mL spectinomycin for unnatural amino acid incorporation). The next day, cells were grown at 37°C in 3 L of buffered terrific broth media with 0.5x antibiotics to OD_600nm_ of 0.8, then induced with 1 mM IPTG at 18°C overnight. For unnatural amino acid incorporation, 2 mM AzF was added per liter before induction with IPTG. Cells were harvested by spinning at 3,500 rpm for 20 minutes at room temperature and then resuspended in lysis buffer supplemented with protease inhibitors (aprotonin, leupeptin, pepstatin, and AEBSF), benzonase, and 2mg/mL lysozyme.

The lid subcomplex was purified in two affinity steps and one size exclusion step. First, cells were lysed by sonication and centrifuged for 30 min at 15,000 rpm at 4°C for lysate clarification. The eluant was loaded to a 5 mL HisTrap FF crude column using a peristaltic pump, washed with NiA buffer, and eluted with NiB buffer. The eluant was then flowed on an amylose resin (NEB), washed with NiA buffer, and eluted with NiA buffer supplemented with 10 mM maltose. After concentrating with a 30 kDa MWCO concentrator, HRV3C protease was added to the amylose eluate for overnight cleavage of the MBP domain at 4°C. After cleavage, the subcomplex was further purified by SEC using a Superose 6 increase 10/300 column in GF buffer supplemented with 500 μM TCEP. For labeling via unnatural amino acid, the MBP cleavage reaction and fluorescent labeling were combined. For this, 150 μM DTNB was added to the concentrated amylose eluate after concentrating and incubated for 10 min on ice before addition of 300 μM LD555 (Lumidyne Technologies), followed by addition of HRV3C protease. The reaction progressed overnight at 4°C and was quenched the next day by incubating with 300 μM free AzF for 5 minutes and 5 mM DTT for 30 minutes at 4°C before SEC. The concentration of lid subcomplex and labeling efficiency were determined by quantification of the 280 nm or dye absorbance using an Agilent UV-Vis Spectrophotometer.

### Purification and biotinylation of 20S core particle

The purification of wild-type or AviTag-20S core was performed as previously described. Briefly, the yeast strain containing the desired 20S core particle construct (Supp. Table 12) was grown in yeast extract, peptone, and dextrose at 30 °C for 3 days. The cells were pelleted, resuspended in core lysis buffer (60 mM HEPES (pH 7.6), 500 mM NaCl, 1 mM EDTA, and 0.2% NP-40) and popcorned into liquid nitrogen, then lysed in a Cryomill 6875D (SPEX SamplePrep). The yeast powder was thawed to room temperature, additional lysis buffer was added, and then clarified by centrifugation for 45 min at 15,000 rpm at 4°C for lysate clarification. The 20S core particle was purified using anti-Flag M2 affinity resin. The eluate was concentrated using an 30 kDa MWCO concentrator, and the concentration was determined by absorbance at 280 nm. After biotinylation through incubation with 25 μM *E. coli* biotin ligase BirA and 100 μM D-biotin in the presence of 10 mM ATP and 10 mM MgCl_2_ overnight at 4°C, the complex was further purified by SEC with a Superose 6 Increase 10/300 column, equilibrated in GF buffer supplemented with 0.5 mM TCEP. The extent of biotinylation was determined in a gel shift assay by incubation with NeutrAvidin. For wild-type core particle, the concentrated Anti-FLAG M2 resin eluate was directly loaded on to SEC and the concentration determined by quantification of the 280nm absorbance using an Agilent UV-Vis Spectrophotometer.

### Purification substrates

The titin I27^V15P^ and titin I27 substrates were purified and labeled as described ^17^. Briefly, *E. coli* BL21-star DE3 cells were transformed with pAM91 or pAM93 (Supp. Table 11). Cells were cultured overnight in DYT media supplemented with 1x antibiotics (50 μg/mL kanamycin). The next day, cells were grown at 30 °C in 3 L of DYT media with 1x antibiotics to an OD_600nm_ of 0.7, then induced with 1 mM IPTG at 30 °C for 3 hours. Cells were harvested by spinning at 3,500 rpm for 20 minutes at room temperature and then resuspended in chitin buffer (60 mM HEPES pH 7.6, 150 mM NaCl, 2 mM EDTA, 5% glycerol) supplemented with protease inhibitors (aprotonin, leupeptin, pepstatin, and 4AEBSF) and benzonase. After lysis and clarification, the lysate was batch bound to a 15 mL of chitin resin (NEB) for 1 hour at 4°C. Then, the resin was washed with 50 mL of chitin buffer supplemented with proteasome inhibitors, 500 mM NaCl, and 0.2% Triton X-100. The substrates were eluted by overnight incubation with chitin elution buffer (60 mM HEPES pH 8.5, 150 mM NaCl, 2 mM EDTA, 5% glycerol, and 50 mM DTT). The eluate was concentrated using a 10 kDa MWCO concentrator and loaded on a HiLoad 16/600 Superdex 75pg column (Cytiva) with GF buffer for SEC.

### Labeling of substrates

Fluorophores were covalently attached to the substrate at its N-terminus for fluorescence polarization or gel-based measurements, or in a single engineered cysteine in the substrate unstructured initiation region for the ensemble or single molecule substrate processing assay. For N-terminal labeling, 100 μM substrate was incubated for 90 minutes at room temperature with 20 mM recombinant SortA enzyme and 500 μM FAM-containing peptide (5-carboxyfluorescein-HHHHHHLPETGG, Biomatik) in GF buffer supplemented with 1 mM DTT and 10 mM CaCl_2_. The labeled substrate was enriched for by clean up using a 1mL HisTrap High Performance (Cytiva), followed by SEC on a Superdex 75 10/300 column (Cytiva) equilibrated with GF buffer. For cysteine labeling using maleimide chemistry, substrates were dialyzed for 3 hours at room temperature into labeling buffer (30 mM HEPES 7.2, 150 mM NaCl, 2 mM EDTA) using a Slide-a-Lyzer mini dialysis cup (ThermoFisher). Then, the substrate was diluted to 100 μM and incubated for 10 minutes at room temperature with 200 μM SulfoCy5 (for stopped-flow measurements, Click Chemistry Tools) or LD555 (for smFRET measurements). Excess dye was quenched with 2 mM DTT and removed by SEC on a Superdex 75 10/300 column (Cytiva) with GF buffer supplemented with 500 μM TCEP. The substrate concentration and labeling efficiency were determined by absorbance measurements using an Agilent UV-Vis Spectrophotometer.

### Purification of Rpn10, ubiquitin and ubiquitination machinery

Yeast Rpn10, *M. musculus* Uba1, Ubc4, Rsp5, and ubiquitin were prepared as described in ^30^ using standard expression and purification procedures ^30,45,46^.

### Substrate ubiquitination

Enzymatic addition of ubiquitin chains to titin I27 substrates containing an unstructured region with a PPPYX motif and a single lysine residue was performed by incubating 10 μM substrate for 50 minutes at 25°C in GF buffer with 10 mM ATP, 200 μM ubiquitin, 2.5 μM *M. musculus* E1, 2.5 μM UbcH7, and 25 μM Rsp5. The extent of substrate ubiquitination was determined by 4-12% Tris-Glycine SDS-PAGE.

### Single Turnover Degradation measured by fluorescence anisotropy

26S proteasomes were reconstituted to 2x concentration with limiting concentrations of 20S core particle (1 μM final) and saturating concentration of base, lid, Rpn10 (2.5 μM each) in GF buffer supplemented with 0.5 mg/mL BSA, 0.5 mM TCEP, 5 mM ATP, and 1x ATP Regeneration System (creatine phosphokinase and creatine phosphate) for 3 min at room temperature, followed by a 3 min incubation at 30°C. Ubiquitinated substrate was prepared to 2x concentration (100 nM final) by diluting in GF buffer supplemented with 0.5 mg/mL BSA and 500 μM TCEP and incubated for 3 min at 30°C. Ubiquitinated (WT or V15P) FAM-labeled titin I27 (5 μL) was rapidly mixed into reconstituted 26S proteasomes (5 μL) loaded in a 384-well clear bottom plate (Corning). Data was acquired at 30°C in a Synergy Neo2 Multi-Mode Plate Reader (Biotek) by following the changes in fluorescence polarization. Gain values were based on the ubiquitinated substrate alone signal. The resulting curves were fit to a double exponential decay to obtain the degradation rates. Reaction endpoint samples were run on an 4-12% Tris-Glycine SDS-PAGE gel to visualize degradation products.

### ATPase assay

ATP-hydrolysis rates were determined using an NADH-coupled assay as described previously (Beckwith et al., 2013). Briefly, 26S proteasomes were reconstituted to 2x concentration under base-limiting conditions with 200 nM base, 800 nM core, 1.2 μM lid, and 1.5 μM Rpn10 in GF buffer supplemented with 0.5 mg/mL BSA, 0.5 mM TCEP, 5 mM ATP and 0.5 mM TCEP at room temperature for 3 min. Then, 2x ATPase mix (final concentrations: 1 mM NADH, 2.5 mM ATP, 7.5 mM phosphoenolpyruvate, 3 U/mL pyruvate kinase, and 3 U/mL lactate dehydrogenase) was rapidly mixed with the proteasomes. The samples were transferred a 384-well clear bottom plate (Corning) and centrifuged at (1000 x g) for 1 min prior to measurement. Steady-state depletion of NADH was assessed by measuring the absorbance at 340 nm in a Synergy Neo2 Multi-Mode Plate Reader (Biotek).

### Ensemble tail insertion measurements

26S proteasomes were reconstituted to 2x concentration with limiting concentrations of SulfoCy3-labeled base (100 nM final) and saturating concentration of core particle (300 nM final), lid (500 nM final), Rpn10 (600 nM final) for in GF buffer with 0.5 mg/mL BSA, 500 μM TCEP, 5 mM ATP, and 1x ATP Regeneration System (creatine phosphokinase and creatine phosphate) for 3 min at room temperature, followed by a 3 min incubation with oPA (3 mM final) at room temperature. SulfoCy5-labeled ubiquitinated titin I27^V15P^ was prepared to 2x concentration (2 μM final) by diluting in GF buffer with 0.5 mg/mL BSA and 500 μM TCEP. Data was acquired at 25°C using an Auto SF-120 stopped flow instrument (Kintek Corporation) by rapidly mixing 26S proteasomes and ubiquitinated substrate. An excitation wavelength of 532 nM was used, and the Cy3- and Cy5-emission channels were monitored simultaneously. Gain values were determined by mixing ubiquitinated substrate with WT base subcomplex, where no FRET is expected but the fluorescence emission of the donor can be tracked. Since there were limiting amounts of FRET-donor (base) and excess of the FRET-acceptor (substrate), the fast phase of the Cy3 fluorescence quenching was fit to a single exponential decay function using Prism9 to obtain the amplitude and tail insertion time (Tau, s). The curves and fits were normalized to the initial fluorescence for plotting.

### Microscope set up

Single-molecule data acquisition was performed at room temperature with using an inverted microscope Nikon Eclipse Ti2 (Supp. Fig.2), built on a ultrastable single molecule stage. The donor (LD555, Lumidyne Technologies) and acceptor (LD655, Lumidyne Technologies) dyes were imaged using a Nikon LUN-F laser box equipped with 532nm and 640nm lasers operated at 5mW and 2mW power, respectively. We used a filter cube from Chroma (TRF59907) placed on the Eclipse Ti2 for imaging. A 60x oil objective (Nikon) with a 1.49 numerical aperture was used for TIRF, which we utilized with Cargille DF immersion oil and perfect focus system (PFS) to provide active feedback stabilization. The emission from donor and acceptor dyes was separated using image splitting optics (W-View Gemini from Hamamatsu) equipped with a T640lpxr-UF2 dichroic mirror (Chroma) and ET595/50-m and ET655lp bandpass filters (Chroma) and simultaneously imaged with an EMCCD camera (Andor Technology, iXon Ultra 512 x 512 DU-897) cooled to -70°C.

For registration of the two emission channels, images were acquired with 100 nm TetraSpeck beads (Thermo Fisher Scientific). Reaction chambers for TetraSpeck beads were assembled on microscope slides using double-sided Scotch tape and glass coverslips. For preparation of the TetraSpeck bead sample, a 1:10 dilution in GF buffer was flowed into reaction chamber assembled with a glass microscope slide using double-sided tape and glass coverslips. The dilution was incubated for 2 min, the excess beads were washed out using GF buffer, and the sample was imaged using the 532 nm laser. Registration was then performed by overlaying cropped regions from the donor and acceptor emission channels such that the bead positions in each channel agreed. The data acquisition for TetraSpeck beads was performed with the following camera settings: EM gain at 4, readout rate at 17 MHz, and preamp set to 3, with activated overlap mode and camera internally triggered through the NIS-Elements software (Nikon).

### Single-molecule fluorescence microscopy data acquisition

Data was acquired with slight modifications to our previous protocol ^38^. Briefly, reaction chambers were assembled on glass microscope slides using double-sided tape and polyethylene glycol (PEG)–coated coverslips with low-density PEG-biotin (MicroSurfaces Inc.). The reaction chambers were incubated with NeutrAvidin (0.25 mg/ml; Thermo Fisher Scientific), and excess NeutrAvidin was removed by washing with 40 μl of assay buffer (GF buffer supplemented with 0.3 μM Rpn10, 1.2 mM Trolox, 0.2 mM β-mercaptoethanol, 0.5 mg/mL BSA, 2 mM ATP, and an ATP regeneration system (creatine kinase 0.03 mg/ml and 16 mM creatine phosphate)). Proteasomes were reconstituted by incubating 500 nM biotinylated core, 400 nM base, 600 nM lid, and 1 μM Rpn10 in assay buffer supplemented with 0.5 μM TCEP for 10 min at room temperature. Reconstituted proteasomes were flowed into the reaction chamber at 1 nM incubated for 5 min. Excess proteasomes in solution were washed out with 40 μl of assay buffer, followed by 20 μl of imaging buffer (assay buffer supplemented with the protocatechuic acid (PCA, Sigma Aldrich)/protocatechuate 3,4-dioxygenase (PCD, Sigma Aldrich) oxygen scavenging system). For the conformational change assay, 0.5 μM or 1 μM unlabeled ubiquitinated substrate was added to the imaging buffer. For the ATPγS samples in the conformational change assay, proteasomes were washed after the 5 min incubation with assay buffer containing 2 mM ATPγS in place of ATP and the ATP Regeneration system, followed with imaging buffer containing the same substitution. For the substrate-processing assay, 100 nM LD555-labeled substrate was added to the imaging buffer.

For all single-molecule acquisitions, the acceptor fluorophores were imaged first by taking a single frame snapshot after a 640 nm laser excitation, which we used to evaluate the extent of surface deposition and localize particles labeled with the LD655 dye. Then, the donor and acceptor fluorescence signals were simultaneously monitored following 532 nm laser excitation. The acquisition rate for these measurements was 20 Hz and the acquisition settings for proteasome samples were done using the same parameters as TetraSpeck beads described above, but with a higher EM gain of 40. The degradation activity of reconstituted, dye- and biotin-labeled proteasomes was confirmed in bulk assays as previously described ^17,38^.

### Single-molecule substrate processing assay data analysis

We analyzed only singly capped and singly labeled proteasomes, which were identified based on their total fluorescence and a single photobleaching step. Data were analyzed with slight modifications compared to our previous study ^38^. Briefly, raw images were aligned a custom MATLAB script (https://github.com/jabard89/tirfexplorer) and loaded on Fiji to parse using the plugin Spot Intensity Analysis (https://imagej.net/plugins/spot-intensity-analysis) to select particles. The generated traces were then sorted and analyzed using a custom-built MATLAB app as previously described ^38^. The start and end of each substrate-processing event were determined from the total fluorescence, which depends on the residence time of a donor-labeled substrate, and the apparent FRET efficiency values outside substrate processing events were not calculated (indicated by dashed lines in all related figures) for easier scoring of substrate processing events. Substrate processing traces were inspected manually and scored for various features to distinguish the processing steps of degradation. The tail insertion phase was measured from the appearance of a donor signal and intermediate apparent FRET efficiency (∼0.50) to the time of the first high–FRET efficiency peak. Deubiquitination was measured as the phase between the first high-FRET efficiency peak and the last peak before donor recovery (FRET-signal decay). The time for the pre-unfolding dwell was measured by measuring the time spent at ∼0.40 FRET efficiency after tail insertion or deubiquitination, until decay from ∼0.20 FRET efficiency. The translocation phase was scored as starting at the FRET-efficiency signal decay to ∼0.20 and with the persistence of the donor signal, and ending when the donor signal was lost. The values obtained for each scored step were plotted as one-cumulative distribution functions (1-CDF) using Prism9.

To determine the substrate-capture success, the fractional number of productive binding events leading to degradation relative to the total number of substrate encounters (both productive and nonproductive) were calculated. Since substrates were donor-labeled and proteasomes acceptor-labeled, we considered only substrate binding events where an increase in donor fluorescence colocalized with acceptor fluorescence, leading to an apparent FRET efficiency value of 0.50 upon substrate binding. Binding events were scored as productive and indicative of degradation if they showed a high-FRET phase followed by an extended donor recovery and fluorescence dwell as a measure for successful threading after unfolding. Binding events were scored as nonproductive if the ∼0.50 FRET efficiency was not followed by an increase to high– FRET efficiency or recovery of the donor fluorescence. A minimum of eight movies were scored, containing at least 22 binding events per movie, were analyzed for each condition, and the error is the SD for the total number of technical replicates.

To determine the substrate-release following the pre-unfolding state in substrate-processing assays, we calculated the fractional number of events where a persistent pre-unfolding dwell was observed relative to the number of events where degradation was completed (productive events). Substrate processing was evaluated as complete if there was a decrease in the FRET efficiency from ∼ 0.40 to ∼ 0.20 (signifying a progression from the pre-unfolding dwell to translocation of the unfolded protein) and then persisting at ∼ 0.20 during translocation. Substrate release was evaluated as the abrupt loss of donor and acceptor signal during the unfolding dwell, i.e. at the ∼ 0.40 FRET-efficiency state, with no decrease in FRET signal from ∼ 0.40 to ∼ 0.20 or a persistent 0.20 FRET efficiency. As a control to account for dye photobleaching, the LD555 donor survival was determined by assembling 26S proteasomes with LD555-labeled base subcomplex, which were prepared and imaged as described above. The survival of the LD555 emission was calculated by measuring the time of photobleaching for single-labeled particles. A total of 152 particles were measured, which yielded 113 photobleaching events that were plotted in a 1-CDF plot.

### Single-molecule conformational dynamics assay data analysis

Proteasome conformational dynamics data were analyzed using new Hidden Markov Model - Markov Chain Monte Carlo (HMM-MCMC) sampling techniques that implement a Bayesian non-parametric framework. Bleed-through noise was removed from movies after collecting images without samples and adjusting the signal using the provided EMCCD camera gain parameters (EM gain 40, readout rate 17 MHz, preamp 3). Particles were selected based on the presence of both donor- and acceptor-signal fluorescence and traces were generated and filtered to obtain proteasomes that showed stable total fluorescence, were singly capped, singly labeled, and showed transitions between FRET-states. For each smFRET time trace, over 20 000 samples for state trajectory, rates, and FRET efficiencies were generated and pooled together in a bivariate probability distributions and transition-density plots. The mean values of the high and low FRET states were selected as the transition rate for each condition and mutant tested. The scripts for kinetic analysis are available through a GitHub repository at https://github.com/LabPresse/BNP-FRET. FRET efficiency distribution plots were generated using the Spartan software, as previously described ^38^. Substrate processing dwells were scored manually from the onset to the stable end of high apparent FRET efficiency, including brief excursions to low–FRET efficiency during the processing of the titin I27^V15P^ substrate.

### Multiple turnover degradation measured on SDS-PAGE

Proteasomes were reconstituted at 2X concentration with limiting concentrations of 20S core particle with 100 nM core, 400 nM base, 800 nM lid, and 1 μM Rpn10 in GF buffer supplemented with 0.5 mg/mL BSA, 0.5 mM TCEP, 5 mM ATP and 1x ATP Regeneration Mix (creatine kinase and creatine phosphate) for 3 min at room temperature. The FAM-labeled titin I27 substrate was prepared at 2X concentration (4 μM) in GF buffer. Gel-based multiple-turnover measurements of FAM-titin I27 degradation were initiated by addition of 5 μL of ubiquitinated substrate to 5 μL of proteasome sample. Aliquots (1.2 μL) were taken after 60 minutes and quenched in 2X SDS-PAGE loading buffer (5 μL). The reaction samples were run on 16.5% Tris-Tricine SDS-PAGE gels (Bio-Rad) and imaged on a Typhoon variable mode scanner (GE Healthcare) for fluorescein fluorescence.

## References

1. Bard, J.A.M. et al. Structure and Function of the 26S Proteasome. Annu Rev Biochem 87, 697–724 (2018).

2. Verma, R. et al. Role of Rpn11 metalloprotease in deubiquitination and degradation by the 26S proteasome. Science 298, 611–5 (2002).

3. Yao, T. & Cohen, R.E. A cryptic protease couples deubiquitination and degradation by the proteasome. Nature 419, 403–407 (2002).

4. Worden, E.J., Padovani, C. & Martin, A. Structure of the Rpn11-Rpn8 dimer reveals mechanisms of substrate deubiquitination during proteasomal degradation. Nature Structural & Molecular Biology 21, 220–227 (2014).

5. Shi, Y. et al. Rpn1 provides adjacent receptor sites for substrate binding and deubiquitination by the proteasome. Science 351, aad9421 (2016).

6. Deveraux, Q., Ustrell, V., Pickart, C. & Rechsteiner, M. A 26 S protease subunit that binds ubiquitin conjugates. The Journal of Biological Chemistry 269, 7059–7061 (1994).

7. Husnjak, K. et al. Proteasome subunit Rpn13 is a novel ubiquitin receptor. Nature 453, 481–488 (2008).

8. Schreiner, P. et al. Ubiquitin docking at the proteasome through a novel pleckstrin-homology domain interaction. Nature 453, 548–552 (2008).

9. Tomko, R.J., Jr., Funakoshi, M., Schneider, K., Wang, J. & Hochstrasser, M. Heterohexameric ring arrangement of the eukaryotic proteasomal ATPases: implications for proteasome structure and assembly. Mol Cell 38, 393–403 (2010).

10. Matyskiela, M.E. & Martin, A. Design principles of a universal protein degradation machine. J Mol Biol 425, 199–213 (2013).

11. Beckwith, R., Estrin, E., Worden, E.J. & Martin, A. Reconstitution of the 26S proteasome reveals functional asymmetries in its AAA+ unfoldase. Nat Struct Mol Biol 20, 1164–72 (2013).

12. Gates, S.N. & Martin, A. Stairway to translocation: AAA+ motor structures reveal the mechanisms of ATP-dependent substrate translocation. Protein Sci (2019).

13. Erales, J., Hoyt, M.A., Troll, F. & Coffino, P. Functional asymmetries of proteasome translocase pore. J Biol Chem 287, 18535–43 (2012).

14. Hinnerwisch, J., Fenton, W.A., Furtak, K.J., Farr, G.W. & Horwich, A.L. Loops in the central channel of ClpA chaperone mediate protein binding, unfolding, and translocation. Cell 121, 1029–41 (2005).

15. Martin, A., Baker, T.A. & Sauer, R.T. Pore loops of the AAA+ ClpX machine grip substrates to drive translocation and unfolding. Nat Struct Mol Biol 15, 1147–51 (2008).

16. Yamada-Inagawa, T., Okuno, T., Karata, K., Yamanaka, K. & Ogura, T. Conserved pore residues in the AAA protease FtsH are important for proteolysis and its coupling to ATP hydrolysis. J Biol Chem 278, 50182–7 (2003).

17. Bard, J.A.M., Bashore, C., Dong, K.C. & Martin, A. The 26S Proteasome Utilizes a Kinetic Gateway to Prioritize Substrate Degradation. Cell 177, 286–298 e15 (2019).

18. Inobe, T., Fishbain, S., Prakash, S. & Matouschek, A. Defining the geometry of the two-component proteasome degron. Nature Chemical Biology 7, 161–167 (2011).

19. Fishbain, S. et al. Sequence composition of disordered regions fine-tunes protein half-life. Nature Structural & Molecular Biology 22, 214–221 (2015).

20. Yu, H., Kago, G., Yellman, C.M. & Matouschek, A. Ubiquitin-like domains can target to the proteasome but proteolysis requires a disordered region. The EMBO Journal 35, 1522–1536 (2016).

21. Matyskiela, M.E., Lander, G.C. & Martin, A. Conformational switching of the 26S proteasome enables substrate degradation. Nat Struct Mol Biol 20, 781–8 (2013).

22. Beck, F. et al. Near-atomic resolution structural model of the yeast 26S proteasome. Proceedings of the National Academy of Sciences of the United States of America 109, 14870–14875 (2012).

23. Lander, G.C. et al. Complete subunit architecture of the proteasome regulatory particle. Nature 482, 186–191 (2012).

24. Unverdorben, P. et al. Deep classification of a large cryo-EM dataset defines the conformational landscape of the 26S proteasome. Proc Natl Acad Sci U S A 111, 5544–9 (2014).

25. Chen, S. et al. Structural basis for dynamic regulation of the human 26S proteasome. Proc Natl Acad Sci U S A 113, 12991–12996 (2016).

26. Wehmer, M. et al. Structural insights into the functional cycle of the ATPase module of the 26S proteasome. Proc Natl Acad Sci U S A 114, 1305–1310 (2017).

27. Sledz, P. et al. Structure of the 26S proteasome with ATP-gammaS bound provides insights into the mechanism of nucleotide-dependent substrate translocation. Proc Natl Acad Sci U S A 110, 7264–9 (2013).

28. Luan, B. et al. Structure of an endogenous yeast 26S proteasome reveals two major conformational states. Proc Natl Acad Sci U S A 113, 2642–7 (2016).

29. Zhu, Y. et al. Structural mechanism for nucleotide-driven remodeling of the AAA-ATPase unfoldase in the activated human 26S proteasome. Nat Commun 9, 1360 (2018).

30. Worden, E.J., Dong, K.C. & Martin, A. An AAA Motor-Driven Mechanical Switch in Rpn11 Controls Deubiquitination at the 26S Proteasome. Molecular Cell 67, 799–811.e8 (2017).

31. de la Pena, A.H., Goodall, E.A., Gates, S.N., Lander, G.C. & Martin, A. Substrate-engaged 26S proteasome structures reveal mechanisms for ATP-hydrolysis-driven translocation. Science 362(2018).

32. Ding, Z. et al. High-resolution cryo-EM structure of the proteasome in complex with ADP-AlFx. Cell Res 27, 373–385 (2017).

33. Dong, Y. et al. Cryo-EM structures and dynamics of substrate-engaged human 26S proteasome. Nature 565, 49–55 (2019).

34. Eisele, M.R. et al. Expanded Coverage of the 26S Proteasome Conformational Landscape Reveals Mechanisms of Peptidase Gating. Cell Rep 24, 1301–1315 e5 (2018).

35. Huang, X., Luan, B., Wu, J. & Shi, Y. An atomic structure of the human 26S proteasome. Nature Structural & Molecular Biology 23, 778–785 (2016).

36. Lasker, K. et al. Molecular architecture of the 26S proteasome holocomplex determined by an integrative approach. Proceedings of the National Academy of Sciences of the United States of America 109, 1380–1387 (2012).

37. Zhang, S. et al. USP14-regulated allostery of the human proteasome by time-resolved cryo-EM. Nature 605, 567–574 (2022).

38. Jonsson, E., Htet, Z.M., Bard, J.A.M., Dong, K.C. & Martin, A. Ubiquitin modulates 26S proteasome conformational dynamics and promotes substrate degradation. Sci Adv 8, eadd9520 (2022).

39. Greene, E.R. et al. Specific lid-base contacts in the 26s proteasome control the conformational switching required for substrate degradation. Elife 8(2019).

40. Zhang, F. et al. Mechanism of substrate unfolding and translocation by the regulatory particle of the proteasome from Methanocaldococcus jannaschii. Mol Cell 34, 485–96 (2009).

41. Rizo, A.N. et al. Structural basis for substrate gripping and translocation by the ClpB AAA+ disaggregase. Nat Commun 10, 2393 (2019).

42. Dong, Y. et al. Cryo-EM structures and dynamics of substrate-engaged human 26S proteasome. Nature (2018).

43. Aubin-Tam, M.E., Olivares, A.O., Sauer, R.T., Baker, T.A. & Lang, M.J. Single-molecule protein unfolding and translocation by an ATP-fueled proteolytic machine. Cell 145, 257–67 (2011).

44. Maillard, R.A. et al. ClpX(P) generates mechanical force to unfold and translocate its protein substrates. Cell 145, 459–69 (2011).

45. Bashore, C. et al. Ubp6 deubiquitinase controls conformational dynamics and substrate degradation of the 26S proteasome. Nature structural & molecular biology 22, 712–9 (2015).

46. Bard, J.A.M. & Martin, A. Recombinant Expression, Unnatural Amino Acid Incorporation, and Site-Specific Labeling of 26S Proteasomal Subcomplexes. Methods Mol Biol 1844, 219–236 (2018).

